# Timed binary WNT activity is sufficient to model the early human neural axis

**DOI:** 10.64898/2026.10.17.759682

**Authors:** J.O.S. Hackland, G. Anand, R. Vijayakumar, X. Wang, V.D. Bocchi, E.S. Rowland, D. Yang, Y. Wu, S. Mehta, A. Zhong, E. Grossi, R.P. Koche, V. Kirigin Callaú, R.M Walsh, A. Evangelisti, Y. Fan, E. Bernstein, R.J. Garippa, L.A. Stamp, M.M. Hao, T. Zhou, L. Studer

## Abstract

The vertebrate nervous system develops into functionally distinct domains along the body axis^1^. Developmental studies implicate multiple signaling pathways in coordinating this process, but the order of specification events, sufficiency of individual signals, and molecular basis of competency remain unclear^2,3^, leaving the field without a unified model of neuraxis patterning. Enteric nervous system (ENS) and midbrain dopaminergic (mDA) progenitors have reached clinical and preclinical testing^4–11^, yet neither is produced at full purity, and yields vary markedly across human pluripotent stem cell (hPSC) lines, reflecting incomplete control of axial identity.

Here we use single-cell genetic fate mapping to demonstrate that hPSCs can commit to anterior or posterior cranial fate within the first 24 hours of differentiation, before neural induction. Building on this finding, we show that the duration of a single saturating WNT pulse is sufficient to specify three “first-order” domains: anterior cranial (0 h), posterior cranial (24 h) and trunk (48 h), each pre-neural and competent for CNS, neural crest and epidermal lineages. Nested “second-order” domains are specified within first-order domains in response to WNT, FGF, and retinoic acid. Mechanistically, an anterior gene regulatory network repressing WNT targets is irreversibly shut down during the transition to posterior identities. Applying this binary WNT patterning logic, we generate mDA neurons and ENS progenitors robustly across hPSC lines and establish an assembloid model combining first-order progenitors to produce all major regions of the nervous system.

## Introduction

The basic inductive principles of primary axis (antero-posterior) patterning were elucidated through clever experimental embryology largely in amphibian models in the mid 20^th^ century. This work demonstrated that separate inductive elements were responsible for neural formation and the acquisition of axial identity. This process was at different times referred to as *evocation/individuation*^12^, *neuralization/mesodermalization*^13^, or *activation/transformation*^14^. In that era, the prevailing theory was that neural induction occurred before axial patterning with anterior neuroectoderm as default. However, anterior versus posterior character could only be identified through the qualitative analysis of neural structures, and not in epidermal ectoderm. With the advent of the molecular age^15^, multiple pathways were identified as being involved in conveying axial identity, including WNT, FGF and Retinoic acid (RA) signaling^16–28^. More recently, the idea that all neural tissue passes through a default anterior neural progenitor has been challenged with strong evidence for a posterior epiblast state that gives rise to neuro-mesodermal progenitors (NMPs; axial progenitors), responsible for the formation of the spinal cord and probably the trunk neural crest^2,6,29–54^. However, whether NMPs present a special case or whether axial identity is generally established before neural commitment, across the whole axis, has not been tested directly.

Cultures of differentiating hPSCs typically behave in ways that mirror developmental pathways, but several lineages remain intractable and even well-established differentiation protocols exhibit a high degree of variability between hPSC lines, between different labs, or when transitioning a protocol to cGMP and large-scale manufacturing as required for cell therapy. Some of the most challenging lineages are developmentally derived from intermediate zones along the primary axis, where differentiation approaches are focused around careful dosing of morphogens rather than application of strong, binary signals that can be used to specify other regions. Two examples are differentiation of hPSCs into the enteric nervous system (ENS)^4–6,55–60^ or into midbrain dopaminergic (mDA) neurons^61–69^. Although these two lineages are developmentally distant from each other, their generation from hPSCs faces complementary issues. Cultures of hPSC-derived vagal neural crest cells (NCCs), the precursor to the ENS, are often contaminated with anterior cranial neural crest, and cultures of mDA progenitors can be easily contaminated with posterior (hindbrain or even spinal cord) lineages^70,71^. In both cases, a partial failure to accurately impart axial identity is central to the appearance of the off-target cell types.

Multiple distinct protocols and developmental rationales have been proposed to improve the efficiency of generating difficult lineages, which leaves the field without an accepted “best practice” approach and without an integrated model for primary axis patterning. Here we apply modern lineage tracing techniques with integrated genomics to redevelop our approach to axial patterning for mDA neurons and vagal NCCs. We find that WNT signaling is sufficient to determine axial identity, that commitment occurs early, before neural induction, and that all neural derivatives arise from one of three distinct regional progenitor states. Based on this logic we establish a unified strategy for deriving the human neural axis from hPSCs.

## Results

### Early acquisition of vagal competence in hPSC-derived neural crest

In the differentiation of hPSC into neural ectoderm-like derivatives, Retinoic Acid (RA) has long been used to posteriorize cultures of cells that would otherwise adopt an anterior identity^60,72–75^. This effect of RA *in vitro* reflects the well-characterized role the morphogen plays in primary axis patterning during vertebrate development^20–23,76^. The posteriorizing action of RA on hPSC-derived neurectoderm has largely been demonstrated through observation of increased expression of vagal markers, as measured using whole-population resolution techniques such as qPCR and bulk RNA sequencing^5,60,72^. We found that assessing vagal identity of hPSC-derived neural crest cells (NCCs) using single-cell resolution approaches revealed that only a subset of cells adopt vagal identity. This is despite transcriptional changes occurring in the whole population in response to RA exposure and without any loss of neural crest induction efficiency (*SOX10::GFP*+ p75++; Fig. 1a-c).

**Figure 1.**
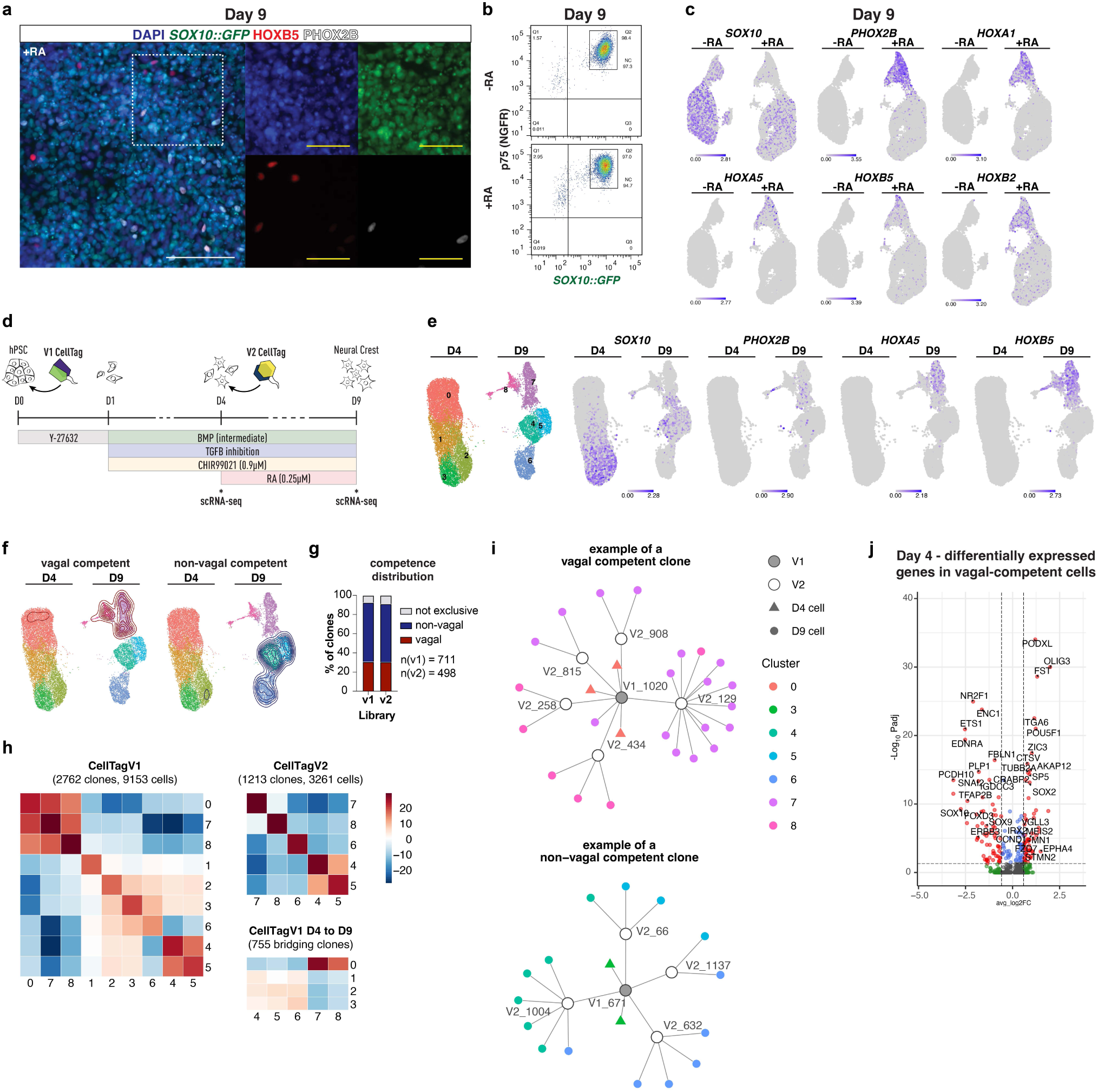
Single cell genetic fate mapping reveals early posterior commitment in vagal neural crest differentiation, **(a)** ICC of day 9 neural crest differentiation cultures using *H9-SOX10::GFP* reporter line in the presence of retinoic acid (RA). *SOX10::GFP* induction indicates neural crest identity in majority of cells. Vagal markers H0XB5 and PH0X2B only detectable in a minority of cells, (b) Flow cytometry using the *H9-SOX10::GFP* reporter and p75 demonstrating no loss of NC induction efficiency in the presence of retinoic acid. This is despite low levels of *SOX 10* transcripts detected in the RA treated condition in (c). (c) scRNA-seq gene expression overlay showing culture-wide transcriptional changes in response to retinoic acid, but only a subset of cells responding with the upregulation of vagal markers *PH0X2B HOXA1/A5/B2/B5.* **(d)** Schematic depicting CellTag experiment using two barcode libraries (applied day 0 & day 4) and two scRNA-seq timepoints, **(e)** UMAPs for CellTag experiment showing clustering and neural crest/vagal marker expression overlay, **(f)** UMAP with density overlay showing location of cells belonging to vagal competent or non-vagal competent clones, **(g)** Proportion of clones showing exclusive bias towards or against vagal competence for each library, **(h)** Clonal coupling analysis showing z-scores quantifying clonal relationships between clusters for CellTagVI and CellTagV2. **(i)** Network diagrams showing representative high cell number vagal competent and non-vagal competent clones, (j) Volcano plot showing enriched/depleted gene expression in vagal competent clones at day 4 (before application of retinoic acid). Scale bar: White - 100pm; yellow - 50*μ*m.

To investigate why some cells adopt a vagal fate while others do not, we applied the genetic fate mapping tool “CellTag”^77^. CellTag uses a lentiviral delivery system to sequentially introduce up to 3 libraries of heritable transcribed barcodes that can be sequenced as part of a routine scRNA-seq pipeline. This allows for the construction of multilevel cell lineage trees alongside a time course of transcriptional identity and thus the identification of developmental commitment or bias in differentiating cells. To avoid rapid silencing of the CellTag construct we switched the original CMV promoter to EF1a (Supp. Fig. 1a). We then applied two CellTag libraries to vagal NCC differentiation cultures at the onset of differentiation (D0; 2hr post-seed; V1) and at the onset of RA exposure (D4; V2; Fig. 1d). Samples were taken for scRNA-seq at day 4 (D4; immediately before RA exposure) and at day 9 (D9; differentiation endpoint; Fig. 1d). Analysis of the data revealed D4 cultures contain both cells with a developmentally mature signature (SOX10+/POU5F1-; clusters 2&3) and a less mature signature (SOX10-/POU5F1+; cluster 0; Fig. 1e, Supp. Fig. 1b). Expression of the ectodermal pioneer factor *TFAP2A* was widespread at this stage (Supp. Fig. 1b). At D9, clusters 4-7 exhibited a gene expression signature consistent with neural crest and neural crest derivatives (expression of *SOX10*, *TFAP2A*, *TWIST1*; Fig. 1e, Supp. Fig. 1b), while cluster 8 appeared to be a small number of off-target CNS derivatives (*PAX6*+; Supp. Fig. 1b). Notably, expression of vagal markers was entirely restricted to a sub population of cells (clusters 7 and 8; *HOXA5*+, *HOXB5*+, *PHOX2B*+; Fig. 1e, Supp. Fig. 1b) as had been seen previously (Fig. 1c) and included both CNS- and neural crest-like cells and derivatives (Supp. Fig. 1b).

Lineage reconstruction using the CellTagR package^77^ and randomized testing to identify lineage-enriched clones revealed that 92% of V1 clones (with 3 or more cells) exhibited complete restriction to either vagal (clusters 7-8) or non-vagal (clusters 4-6) fates (Fig. 1g). This was supported by a similar pattern of commitment in V2 clones established 4 days later. Clonal coupling analysis on V1 clones (with 2 cells or more) demonstrated a high degree of co-occupancy across clusters 0, 7-8 or 1-6 (Fig. 1h). The same approach with V2 clones showed a greater degree of segregation, indicating that cells were largely already committed to cluster (with exception of c4/5; Fig. 1h). Together these data indicate that most cells commit toward or against vagal competence within one cell cycle (∼24hr) from differentiation onset. This point is demonstrated by the UMAP location of cells sequenced at D9 that belong to fate-restricted clones (Fig. 1f, Supp. Fig. 1d) and by network diagrams of representative high cell number clones from each group (Fig. 1i).

Intriguingly, cells from committed clones at D4, before the onset of RA exposure, also exhibited a distinct transcriptional profile between vagal competent and non-vagal competent groups. Cells belonging to vagal competent lineages were largely in cluster 0, while cells belonging to non-vagal competent lineages were most concentrated in cluster 2 (Supp. Fig. 1d). Gene expression analysis identified several hindbrain-associated genes significantly enriched in the vagal competent clones, including *FST*, *EPHA4*, *MEIS2*, *STMN2* and *VGLL3* (Fig. 1j; Supp. Fig. 1e). These cells also had a developmentally immature transcriptional profile, with genes such as *POU5F1*, *SOX2* and *PODXL* enriched and NCC markers such as *SOX10*, *TFAP2B* and *ETS-1* depleted (Fig. 1j). This indicates that an early, spontaneous, acquisition of posterior identity occurs in a subset of cells, and that these cells respond to RA with upregulation of vagal genes. Two key components of the WNT signaling pathway were also enriched in the vagal-competent cells at D4: *FZD7* and *SP5*. With the implication that WNT signaling might be involved in the acquisition of vagal competence we inhibited the pathway with 5µM XAV939 and observed a reduction in *HOXB5* expression at D9 (Supp. Fig. 1f).

### Transient WNT activation induces vagal competence

With the insight that acquisition of vagal competence occurs very early, we sought culture conditions that would induce this phenotype. Early, strong activation of WNT signaling has been shown to induce a trunk-like identity in differentiating pluripotent stem cells in both the human and mouse (neuro-mesodermal progenitors; NMPs; axial progenitors)^6,34–37,39,41,42,44,46^. In the case of hPSCs, approximately 48hr of activation is required to generate trunk derivatives. Given that vagal competence appears to be established in half this time (based on point of V1 CellTag transduction and estimated cell cycle time), we hypothesized that 24hr of WNT activation may be sufficient for inducing a less posterior, vagal-competent fate. Redesigning our protocol with this in mind and using the small molecule CHIR99021 (CHIR) to mimic canonical WNT signaling resulted in a dramatic increase in vagal marker expression in hPSC-derived NCCs at day 7 (D7; 4.5µM CHIR D0-1 referred to hereafter as WNT “bump”; Fig. 2a-c). This was confirmed across 3 separate hPSC lines (Fig. 2a-b; Supp. Fig. 2a-c). scRNA-seq analysis at day 4 (D4) did not reveal widespread HOX gene expression in the “bump” condition but instead showed enrichment for the retinoic acid binding protein *CRABP1*, as well as hindbrain markers *DACH1*/*DACH2* (Fig. 2d). Some enriched markers were also hits in the vagal-committed clones of the CellTag experiment (*EPHA4*, *FZD7*), while others were significantly enriched in cluster 0 from the CellTag experiment (*CRABP1*, *DACH1*; Supp. Fig. 1. e). Additional increase of WNT activity at later stages (1.5µM CHIR; D4-7) did not further improve vagal induction efficiency, but did improve cell yield (Fig. 2b’, Supp. Fig. 2a-b).

**Figure 2.**
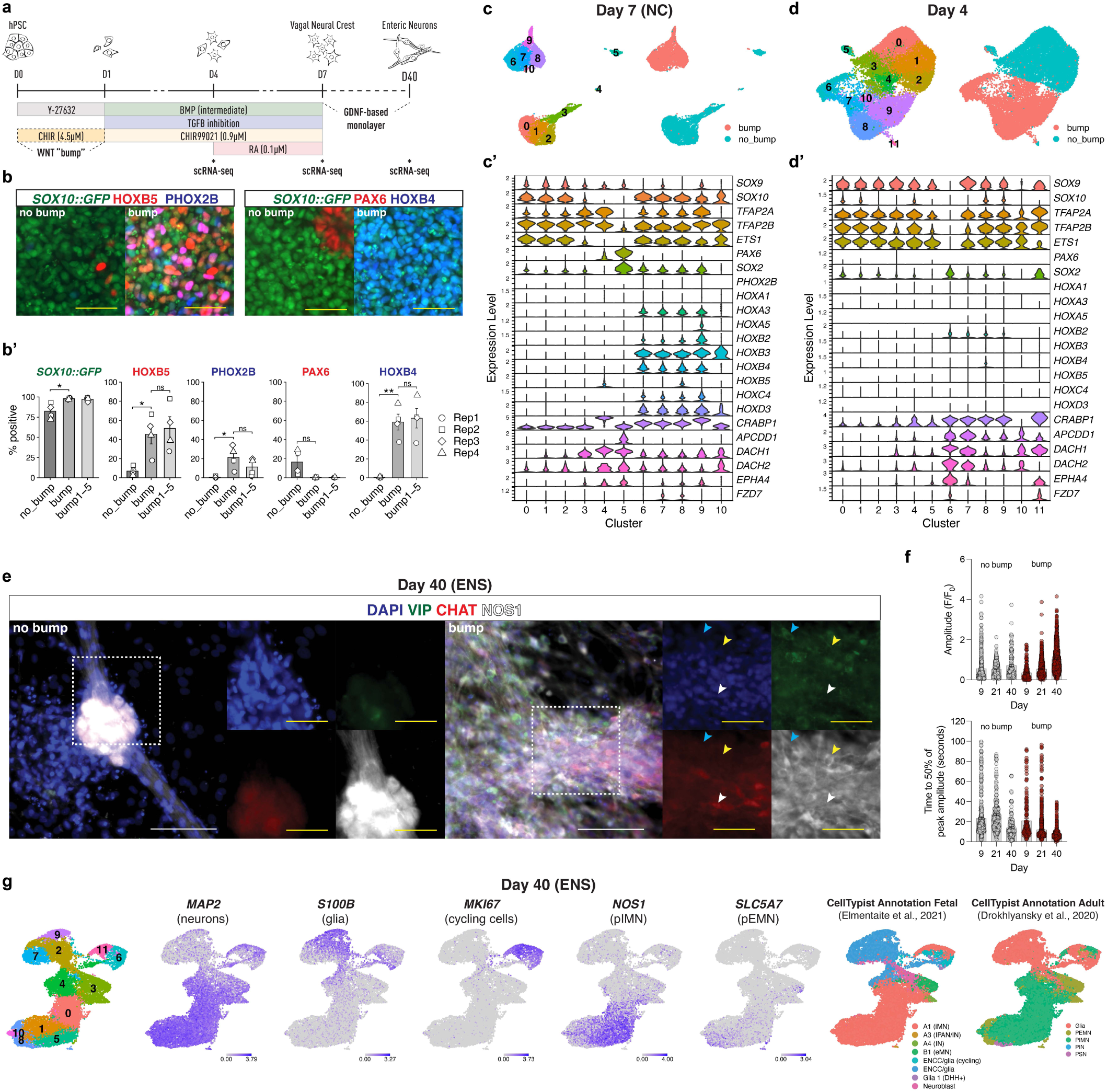
Early, strong, transient WNT activation induces vagal competence in differentiating hPSCs. **(a)** Schematic representing the WNT “bump” protocol. 4.5pM CHIR D0-1 induces a culture-wide shift to vagal competence. ENS differentiation is used to functionally assess identity. Timepoints for scRNA-seq marked, **(b)** ICC (using H9-*SOX 10::GFP)* demonstrating upregulation of vagal markers HOXB5, PHOX2B and HOXB4 at day 7 of NC induction using the WNT bump protocol. CNS marker PAX6 identifies rare contaminating off-target cells in the “no bump” condition. These are largely absent in the “bump” condition, **(b’)** Quantification of ICC data in (b). “bump1-5” is additional trial condition with 1 ,5/vM CHIR D4-7 for which a representative image is not shown, **(c-d)** UMAPS representing scRNA-seq analysis at day 7 (c) and day 4 (d) of the WNT bump protocol. Violin plots showing expression of NC/CNS/posterior markers across clusters in (c) and (d) are presented in (c’) and (d’). (e) ENS D40 ICC using H9 showing VIP, CHAT and NOS detection. Cells can be identified that correspond to putative excitatory motor neurons (CHAT+; white arrow), putative inhibitory motor neurons (NOS1+, VIP+; yellow arrow) and putative secreto-vasodilator neurons (VIP+; blue arrow). Condition inset in white, (f) Summary of amplitude and duration (time to 50% of peak) of [Ca^2+^]_i_ transients in “no bump”- (gray) and “bump”- (red) derived ENS cultures over time, (g) UMAPS representing D40 scRNA-seq clustering, gene expression, and analysis with CellTypist annotation (Fetal: iMN, inhibitory motor neuron; IPAN, intrinsic primary afferent neuron; IN, interneuron; eMN, excitatory motor neuron; ENCC, enteric neural crest cell. Adult: PEMN, putative excitatory motor neuron; PIMN, putative inhibitory motor neuron; PIN, putative interneuron; PSN, putative sensory neuron). ICC quantification data in (b’) are presented as mean ± s.e.m. Each point represents a biological replicate. Two-tailed unpaired t-tests were used with Welch’s correction for unequal variances. *P < 0.05, **P < 0.01, ***P < 0.001; ns, not significant (P ≥ 0.05). Scale bar: White - WO*μ*m; yellow - 50*μ*m.

Vagal NCCs function as the primary origin of the enteric nervous system (ENS). We have previously shown that vagal NCCs generated from hPSCs through the application of RA are capable of differentiating into enteric neuron-like cells and rescuing mouse models of Hirschsprung’s disease^5,6^. However, previous protocols yield large numbers of off-target ectomesenchyme-like cells. Using the new WNT “bump” protocol, we were able to generate high purity ENS cultures with minimal ectomesenchymal contamination by day 40 (D40), unlike “no bump” controls (Fig. 2e-g; Supp. Fig. 2d, 3a-k). Using ICC analysis, it is possible to identify putative excitatory motor neurons (CHAT+/NOS low/VIP-; white arrow), putative inhibitory motor neurons (CHAT-/NOS high/VIP+; yellow arrow) and putative secreto-vasodilator neurons (CHAT-/NOS low/VIP+; cyan arrow; Fig. 2e)^78–83^. Accurate quantification of these thick cultures is difficult, particularly for the “no bump” condition. Therefore, we used scRNA-seq for a more in-depth analysis. Using a combination of Accutase and Papain for dissociation, our analysis revealed distinct *SOX10::GFP*/p75 profiles by flow cytometry, as well as cellular morphology in suspension (Supp. Fig. 3a-b). D40 “bump”-derived ENS cultures contained a majority of neurons (65.4%; *MAP2*+/*TUBB3*+/*PRPH*+), as well as a substantial population of glia (22.9%; *S100B*+/*SOX10*+/*PLP1*+) and cycling progenitors (9.7%; *MKI67*+; Fig2. g; Supp. Fig. 3c-f). Only 2% of cells sequenced corresponded to an ectomesenchymal phenotype (*COL1A1*+; Supp. Fig. 3d). As with the ICC analysis we observed a high proportion of NOS1^high^ inhibitory motor neuron-like cells (Fig. 2g)^80^ and markers for cholinergic neurons could also be detected in two otherwise transcriptionally distinct sets of cells, indicating the presence of excitatory motor neuron-like cells (c3,8; Fig. 2g)^80^. We used the unbiased annotation tool CellTypist^84^ to annotate our dataset using publicly available human data from both the adult^80,81^ and the fetal^81^ ENS (Fig. 2g). In both cases the majority of neurons were identified as putative inhibitory motor neurons while putative excitatory motor neurons, interneurons and sensory neurons were also labeled (Fig. 2g). Cells sequenced at D40 derived from the “no bump” control included a large number of fibroblast-like profiles (*COL1A1*+), although smaller numbers of neurons and glia were also present (Supp. Fig. 3c-e). Treatment with known ENS morphogens EDN3 or GDNF during the second half of the “WNT bump” protocol minimized RA-based toxicity without altering the transcriptional landscape at D40 (Supp. Fig. 3c-e). To further characterize our “bump”-derived ENS cultures we used live cell calcium imaging, an important tool in the assessment of function and maturation in both primary and hPSC-derived ENS cultures^56,85^. We found that when depolarized with extracellular high K^+^, “bump”-derived ENS cultures showed an increased transient amplitude and decreased duration (time to 50% of peak amplitude) over time, indicating rapid neuronal maturation by D40 (Fig. 2f, Supp. Fig. 3i). This occurred more rapidly than in the “no bump” condition and compared to other studies on hPSC-derived ENS^56^. This could indicate that the presence of off-target mesenchymal cells negatively impacts neural maturation, or that the improved early patterning has a cell-intrinsic maturation benefit. We also detected a robust response to the nicotinic acetylcholine receptor agonist DMPP, the NK1 receptor agonist Substance P, and serotonin (5-HT; Supp. Fig. 3j). Finally, we found that “bump”-derived ENS cultures exhibited spontaneous firing activity that initiated in a synchronous manner (day 21) and then developed into asynchronous subnetworks, further indicating maturation (Supp. Fig. 3h)^56,86^.

We also tried alternative posteriorizing factors during differentiation. Treatment with RA between D0-1 (instead of the WNT “bump”) induced a non-neural ectoderm phenotype, which could be compensated for either with 2.0µM DMH1 (D0-1) or 4.0µM DMH1 and 1.2µM CHIR (D1-7; Supp. Fig. 4a). However, in each case this did not result in expression of vagal markers *HOXB5* and *PHOX2B* in the resulting NCCs. Treatment with FGF8 between D0-1 also did not increase the proportion of vagal NCCs (Supp. Fig. 4b). Similarly, treatment with FGF8 between D1-4 or D1-7 in the “no bump” condition did not result in induction of vagal fate (Supp. Fig. 4c). In line with this, treatment with FGF8 (D1-4 or D1-7) in WNT “bump” cultures appeared to block vagal identity (Supp. Fig. 4c).

### Limited induction window for vagal competent cells

The hindbrain-like gene expression signature of vagal-competent clones at D4 of the CellTag experiment led us to hypothesize that the D1 vagal competent progenitors may reflect an early, regionally committed posterior progenitor. The small proportion of *GBX2* positive cells sequenced at D4 in clusters 0-1 is further evidence in support of this hypothesis (Supp. Fig. 1e). *GBX2* is known to be necessary for neural crest formation at the hindbrain level, where it is expressed at the neural plate border but not in migrating neural crest ^87–89^. It therefore could have been more widely expressed in vagal competent cells earlier during differentiation. To study the expression of *GBX2* during vagal neural crest differentiation, we generated a reporter cell line using a tdTomato gene fused to H2B for nuclear localization, as has been previously described^90^ and modulated WNT activation during the first few days of differentiation (Fig. 3a). Using this reporter, we observed that a high level of *GBX2::tdTOM* expression in WNT “bump” cultures at D1 (Fig. 3b; 24hr). This was also detectable at D2 where ICC shows widespread evidence of *OTX2* expression in “no bump” controls but no OTX2 in the WNT “bump” condition (Fig. 3c). A small proportion of *GBX2::tdTOM* positive cells could be found in the “no bump” condition, presumably corresponding to the spontaneous vagal competent population identified in the CellTag experiment (Fig. 3b-c). As expected from our previous studies and those of others ^6,34,35,37,39,41,42,44,46^ extending the treatment with CHIR99021 to a total of 48hr (“long bump”) resulted in induction of trunk markers TBXT (brachyury) and CDX2 (Fig. 3c). The “long bump” condition does not result in vagal competency as is demonstrated by the absence of *HOXB5* expression at D7 in this condition (Fig. 3d, row 1).

**Figure 3.**
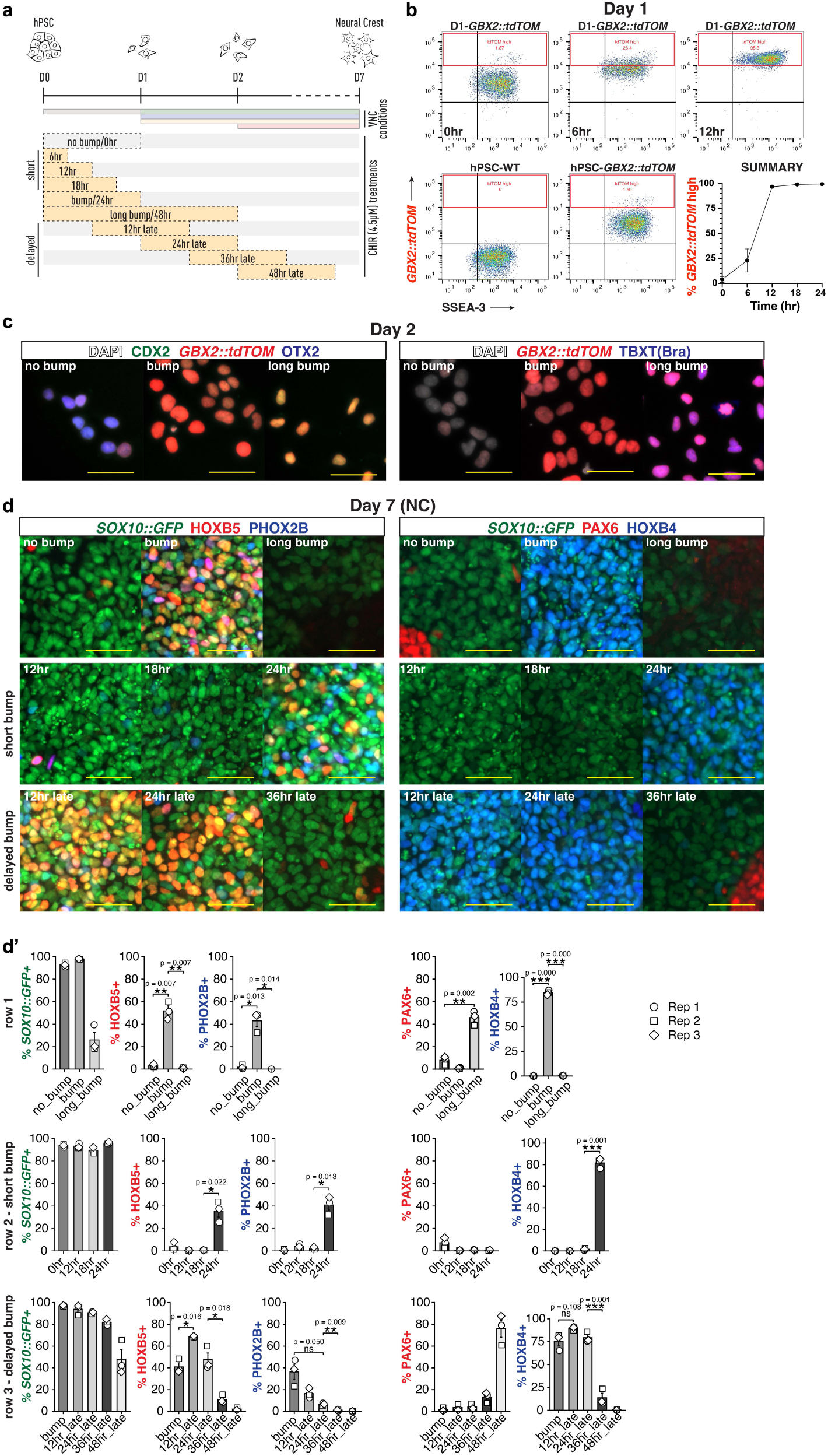
Modulation of early WNT dynamics prior to neural crest differentiation reveals an induction window for vagal competence, **(a)** Schematic showing experimental conditions used to study the role of WNT dynamics in progenitor specification and subsequent vagal neural crest induction, **(b)** Flow cytometry showing population-wide induction of *GBX2* expression at day 1 after just 12 hours of treatment with 4.5pM CHIR. (c) Day 2 ICC showing loss of OTX2 and gain of *GBX2::td-TOM* after 24hr treatment with 4.5pM CHIR (”bump”). CDX2 is gained after 48hr treatment with 4.5pM CHIR (’’long bump”). Also shown is gain of TBXT (Brachyury) only with 48hr of WNT activation (“long bump”), (d) ICC showing that extended 4.5pM CHIR treatment (’’long bump”) results in loss of vagal competency - no HOXB5 at D7 (1st row). Short duration treatment with 4.5pM CHIR (”12hr”, “18hr”) results in loss of vagal competency despite *GBX2*expression seen in (b) (2nd row). A delay in CHIR treatment of more than 24hr results in failure to induce vagal competence. Quantification in (d’). ICC quantification data in (d’) are presented as mean ± s.e.m. Each point represents a biological replicate. Two-tailed unpaired t-tests were used with Welch’s correction for unequal variances. *P < 0.05, **P < 0.01, ***P < 0.001; ns, not significant (P ≥ 0.05). Scale bar - 50μm.

Interestingly, we found that just 12hr of WNT activation was required to induce a population-wide transition to a *GBX2::tdTOM* positive state (Fig. 3b), but a full 24hr WNT activation was required for vagal competence (Fig. 3d, row 2). Delaying CHIR by 36 hours or more resulted in a failure to induce vagal competence (Fig. 3d, row 3). Together these results reveal a window within which cells respond to WNT with vagal competence acquisition.

To test the pan-ectodermal potential of WNT “bump”-induced posterior progenitors we subjected them to dual SMAD inhibition (DSi). As expected, we found that PAX6 positive CNS-like cultures were formed from “no bump”, “bump” and “long bump” cultures (Supp. Fig. 5a). Furthermore, we found that epidermal-like cells could be derived when WNT “bump” progenitors were exposed to BMP4 (15ng/mL; Supp. Fig. 5a). However, these cultures were distinct from those derived from the “no bump” condition, with only a subset of cells positive for TFAP2A despite epidermal-like morphology. The distinct TFAP2A expression might reflect differences in gene expression in the epidermis along the AP axis (as seen with HOX genes)^91^, or a transition to amniotic ectoderm-like derivatives possibly due to culture density ^92^.

### Vagal competent state governed by OTX2-GBX2 antagonism and WNT response elements

We next set out to molecularly characterize the vagal competent state using scRNA-seq analysis in combination with bulk ATAC-seq and CUT&RUN at day 1 (D1; Fig. 4a). D1 “bump” cultures showed widespread activation of *GBX2* and hindbrain-level HOX with a dramatic decrease in *OTX2* transcripts (Fig. 4b). Transcriptional activation of putative vagal coordinator *CDX1* was detected in D1 WNT “bump” cultures as well as “long bump” cultures (Fig. 4b)^93^. Differential gene expression analysis also identified several hits from the CellTag experiment as being significantly enriched in the WNT “bump” condition, including *FST*, *MEIS2* and *STMN2* (Supp. Fig. 6a-b).

**Figure 4.**
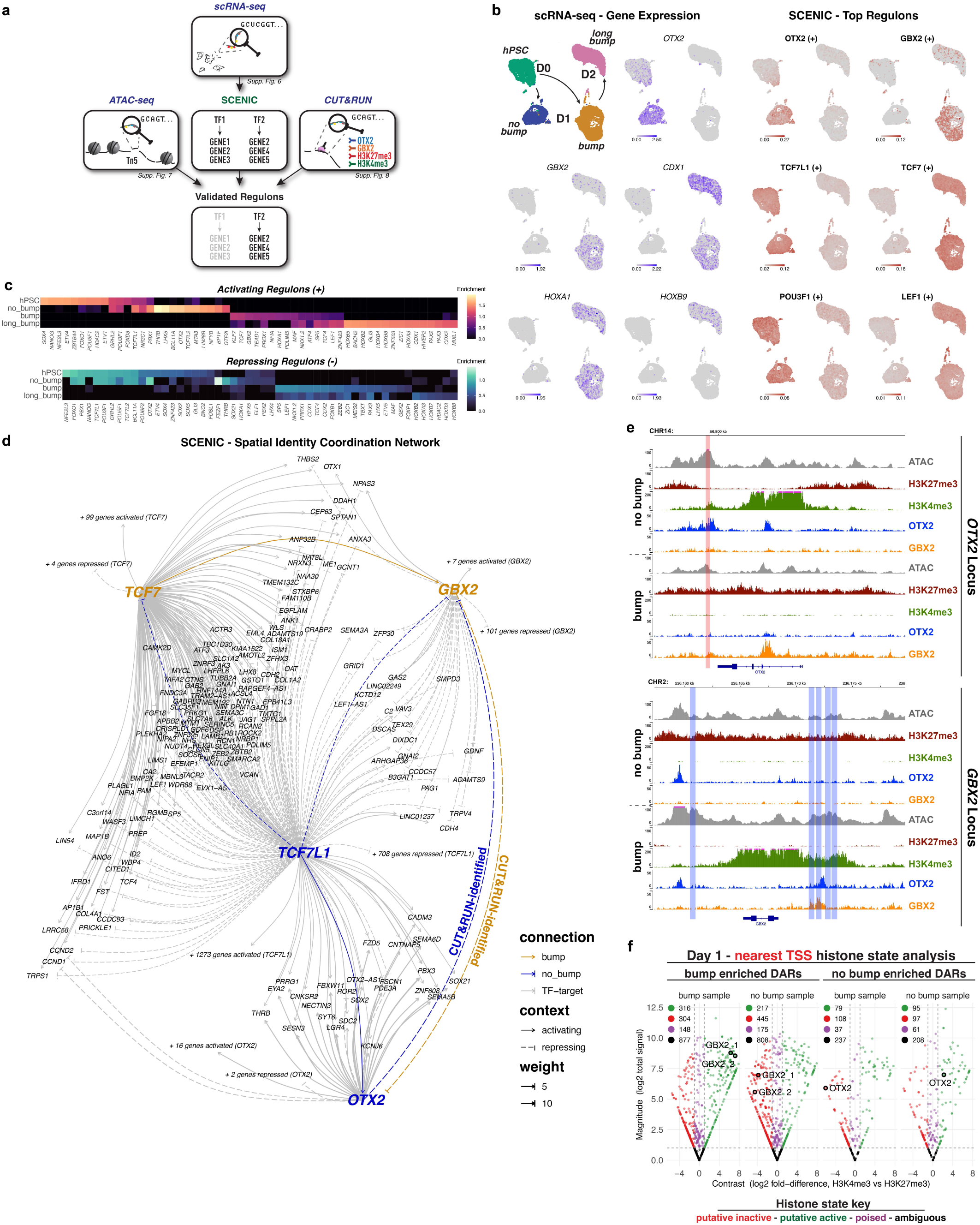
Transition to vagal competence is coordinated by OTX2, GBX2 and WNT response elements, **(a)** Analytical strategy diagram - scRNA-seq data was analyzed using SCENIC to identify a putative gene regulatory network. The output regulons were validated using bulk ATAC-seq and CUT&RUN. (b) scRNA-seq data - UMAP dimensionality reduction with overlays for gene expression of key primary axis markers (columns 1-2) and SCENIC regulon enrichment score for top activating regulons (columns 3-4). (c) Heatmaps showing top 15 activating regulons and top 20 repressing regulons for each condition. Where there is redundancy between conditions regulons are not shown twice, **(d)** Network diagram showing interactions and shared targets between top regulons. Unshared targets summarised under a single edge. No interaction was predicted by SCENIC between OTX2 and GBX2, but mutual binding to loci was identified in the CUT&RUN analysis and represented on this diagram, **(e)** ATAC-seq and CUT&RUN results for the *OTX2* and *GBX2* loci. Vertical bars indicate significant differentially accessible regions (DARs) enriched in no bump (red) and bump (blue), **(f)** Histone state analysis summarizing H3K4me3 and H3K27me3 CUT&RUN data at the nearest TSS to enriched DARs from the ATAC-seq data. Annotated as putative active (high H3K4me3; green; right of plot), inactive (high H3K27me3; red; left of plot) poised (high for both marks; purple; middle of plot) or ambiguous (low for both marks; black; bottom of plot). Total number of TSSs on each plot is recorded at the top left of the plot. Highlighted are TSSs for GBX2 (2x TSS; adjacent to ”bump” enriched DARs) and OTX2 (adjacent to ”no bump” enriched DAR).

Further analysis of these scRNA-seq data using the gene regulatory network (GRN) prediction package SCENIC^94,95^ identified putative activating and repressing regulons involved in the interconnected coordination of cell state across the 4 conditions (Fig. 4b-c; Supp. Fig. 6c). Each of these regulons consists of transcription factor (TF)-target relationships identified via coexpression dynamics and then filtered according to TF consensus sequence abundance and overall regulon enrichment score across the dataset. Both activating and repressing regulons associated with GBX2 and OTX2 were top hits for the WNT “bump” and “no bump” conditions respectively, reflecting their known role as both transcriptional activators and repressors in a target-specific context^96–101^ (Fig. 4b-c; Supp. Fig. 6c). Other top regulons included components of the WNT signaling pathway such as *TCF7*, *LEF1* and *SP5* in the WNT “bump” condition and *TCF7L1* and *TCF7L2* in the “no bump” condition (Fig. 4b-c; Supp. Fig. 6c). These results suggest the “no bump” and “hPSC” states are maintained in part through the TLE/Groucho-mediated suppression of WNT targets by TCF7L1 (and likely TCF7L2). When WNT signaling is activated, TCF7 and LEF1 replace TCF7L1 and function as potent activators when bound to B-catenin, synergizing with SP5 to induce downstream WNT targets. This is a process that has been well described as the core regulatory cascade that coordinates the acquisition of posterior identity *in vivo*, suggesting that this protocol is recapitulating the mechanism of primary axis patterning that occurs during early embryonic development^102–107^. Supporting this are the opposing relationships predicted for targets of TCF7L1 and TCF7 (Fig. 4d) and TCF7L1 and LEF1 (Supp. Fig. 6d). There are also activating relationships predicted by SCENIC for TCF7L1, but these are likely instances where a gene is targeted by TCF7 or LEF1 for repression in “bump” or “long bump” conditions, as this would generate an inverse expression relationship with TCF7L1 and these factors share the TCF family consensus sequence.

The interpretation that the vagal competent state represents an early transition to posterior identity is further supported by the identity of genes near differentially accessible regions (DARs) identified using ATAC-seq analysis. These included known posterior markers that overlapped with the SCENIC analysis and gene expression data (e.g. *GBX2*, *HOXA1*, *CDX1/2*), and when subjected to Gene Ontology (GO) analysis resulted in axial patterning-related terms (Supp. Fig. 7a-c). Results of TF foot printing analysis of these DARs using TOBIAS ^108^ also supported much of the GRN predicted by SCENIC (Supp. Fig. 7d). It is important to note that TOBIAS cannot accurately distinguish between TFs with similar consensus sequences and with the settings used (motif-pvalue = 1e^-4^) the TCF7 family and CDX1/CDX2/HOXA9 cannot be resolved (Supp. Fig. 7e). This explains why TCF7L1/2 are identified as hits in the “bump” condition (rather than “no bump”) and HOXA9 is also identified as a hit in “bump” despite not being expressed.

OTX2 and GBX2 are key regulators of primary axis identity, therefore we performed CUT&RUN analysis to identify their targets. We found that a majority (∼70%) of the “bump” enriched DARs were bound by either OTX2 or GBX2 or both in the WNT “bump” condition, whereas “no bump” enriched DARs are largely occupied by OTX2 (Supp. Fig. 8a). We detected each factor at the locus of the other, supporting a direct, mutually repressive relationship (Fig. 4d-e). Identification of differentially occupied regions using DiffBind identified *OTX2* as the top hit for GBX2, as well as for itself, however *GBX2* was not enriched as a hit for OTX2 despite an apparent peak 5kb upstream of the TSS (Fig. 4e, Supp. Fig. 8b). This is likely due to low overall signal in one of our replicates for GBX2, resulting in a high degree of variance between the two replicates and limiting our number of significant hits. Targets identified in both the SCENIC and CUT&RUN analyses included several that were shared for OTX2 and GBX2, including *TLE2*, *OTX1*, *DIXDC1* and *PAX7* (Supp. Fig. 6e).

As adjacency to an enriched DAR does not always mean a gene is primed for expression, we also performed CUT&RUN analysis for the histone marks H3K4me3 and H3K27me3, associated with active and inactive genes respectively (Supp. Fig. 7a). We found the pattern of enrichment of these signatures supported previous observations of distinct axial identity between our conditions (Supp. Fig. 8c). To summarize the relationship of H3K4me3 and H3K27me3 across DARs, we developed a simple “histone state analysis” whereby normalized signal *magnitude* for each mark is plotted against normalized signal *contrast* (Fig. 4f; Supp. Fig 8d). This approach highlights that DARs are not always associated with an active signature but instead exhibit a range of states. TSSs for *GBX2* are in a putative active state in the “bump” sample, but a putative poised state in the “no bump” sample (Fig. 4f). This is in line with observed *GBX2* expression in the “bump” condition (Fig. 3b-c, 4b, 5c-f) and the 36hr window of retained potential for vagal competent state induction in the “no bump” sample (see “delayed bump” condition in Fig. 3d). In contrast the TSS for *OTX2* is in a putative poised state in the “no bump” condition, but a putative inactive state in the “bump” condition, where it appears the cultures have lost the ability to express *OTX2* (no return to an OTX2 positive state at D2 after CHIR treatment D0-1; Fig. 3c). We found further interesting patterns in WNT signaling pathway and anterior-posterior patterning genes (Supp. Fig. 8d).

### Evidence of specific rhombomere identity after dual SMAD inhibition

Our data indicate that the vagal competent state is in fact a posterior progenitor with axial potential broader than just the vagal neural crest region. To further test this hypothesis, we subjected D1 WNT “bump” cultures to dual SMAD inhibition (DSi) to look for further evidence of hindbrain-level lineage identity. The early hindbrain is divided into 8 discrete domains called rhombomeres that each express a distinct cohort of markers that are highly conserved across species^109–121^ (Fig. 5c). Using these markers, it is possible to get an indication of rhombomeric identity without differentiating cells into terminal neural lineages. As we already had evidence of regional rhombomere 7-8 (r7-8)-like identity derived using the WNT “bump” approach (i.e. vagal neural crest; Fig. 2-3), we next identified conditions capable of generating r1-6-like identity. To this end, we investigated both shortened WNT treatment within the first 24 hours, as done previously for NCC differentiation (Fig. 5a-b), as well as manipulation of WNT, FGF or RA signaling conditions from D1-4 (Fig. 5d-h). To our surprise we found that, although virtually 100% of cells exhibited high *GBX2::tdTOM* signal at D1 after 12hr of WNT activation (Fig. 3b), by D4 in DSi conditions a high proportion of these cells had retained an OTX2 positive state (Fig. 5b). This suggests that despite *GBX2* induction, 12hr of WNT activation is not sufficient to fully engage the posterior GRN identified in Figure 4 in all cells. However, 18hr WNT activation resulted in stable *GBX2::tdTOM* induction until D4 and loss of OTX2 in most cells (Fig. 5b). This is notable given 18hr is not long enough to convey vagal competence (Fig. 3d), suggesting that *GBX2* expression is required but not sufficient for this state. The midbrain-hindbrain boundary (MHB) is a major source of FGF8 and has been implicated in the establishment of both midbrain and hindbrain fate^122–125^. We speculated that treatment with FGF8 may induce an anterior hindbrain identity (e.g. r1), however, no induction of EN1 was observed in this condition for any bump duration. We did observe an increase in expression of *GBX2::tdTOM* in the “no bump” cultures, but this occurred without a significant decrease in the number of OTX2 positive cells and was the only condition where we observed OTX2 positive cells with high levels of *GBX2::tdTOM* (Fig. 5e). Although GBX2 largely marks posterior tissues, this result is compatible with an early (∼ E10.5 in mouse) anterior GBX2+ domain that emerges adjacent to a source of FGF8 in the dorsal thalamus^126,127^.

**Figure 5.**
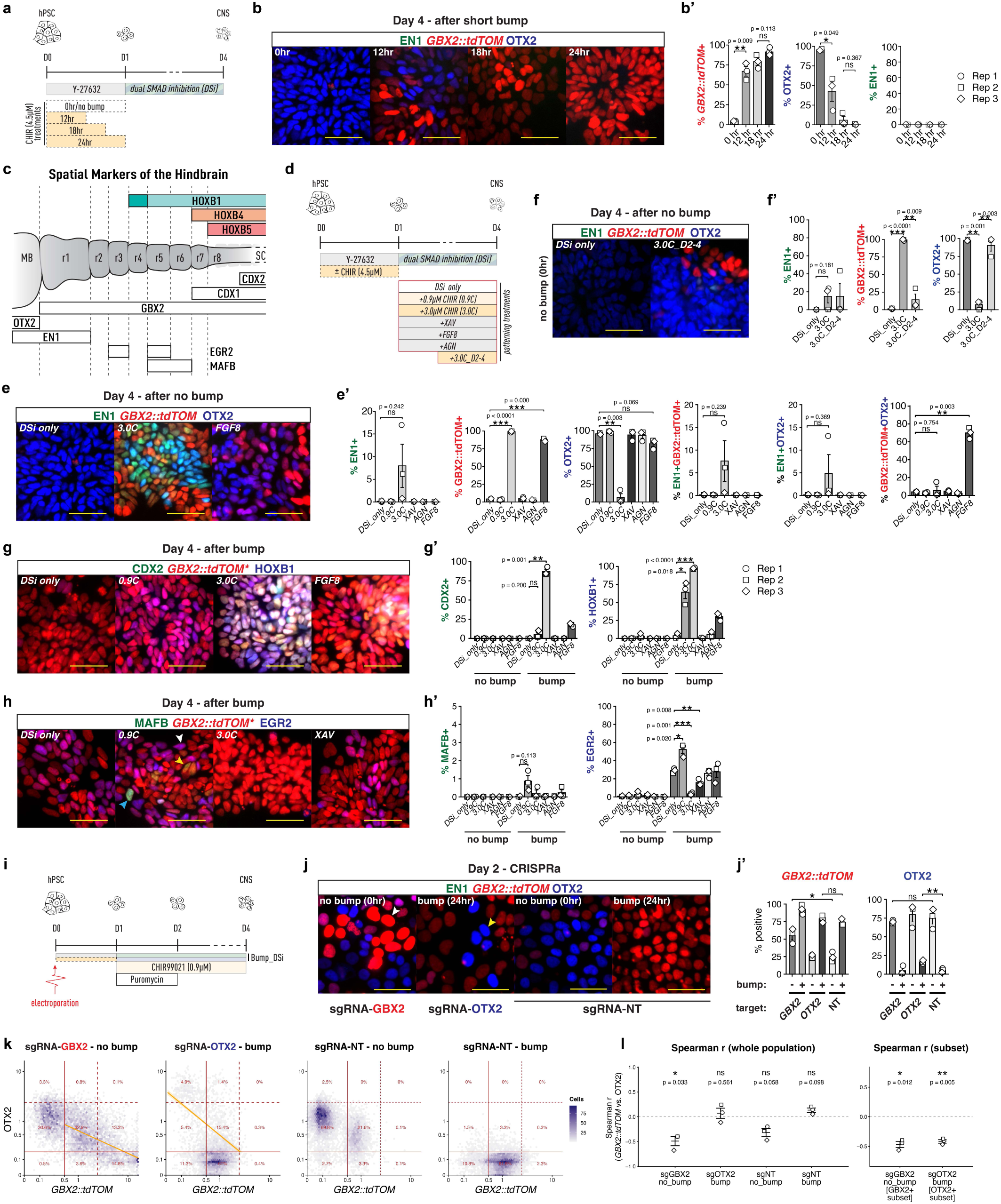
Cells with specific rhombomere-like expression patterns can be generated from day 1 posterior progenitors and OTX2/GBX2 exhibit a functionally antagonistic relationship, **(a)** Schematic showing experimental conditions used in (b). **(b)** Representative day 4 ICC data showing EN1, *GBX2::tdTOM* and OTX2 detection in response to short duration treatment with 4.5;/M CHIR (”12hr”, “18hr”; white inset) followed by dual SMAD inhibition (DSi). Quantification presented in (c’). **(c)** Schematic showing the location of regional markers within the hindbrain. MB = midbrain; SC = spinal cord; r1 -8 = rhombomeres 1 -8. **(d)** Schematic showing experimental conditions used in (e-h). **(e-h)** Representative ICC data for “no bump” or “bump” progenitor cultures subsequently treated with the patterning factors described in (d). Examples of key conditions are presented with the condition inset in white. Arrowheads in (h) represent r3, r5 and r6 cells (white, cyan and yellow respectively). Quantification presented for all conditions tested in (e’-h’). **(i)** Schematic showing design of CRISPRa experiments in dual SMAD inhibition (DSi) conditions with 0.9uM CHIR. **(j)** Representative ICC on day 2 of experiment in (i) with quantification presented in (j’). Arrowheads mark cells with strong activation of *GBX2::tdTOM*and absence of OTX2 (white) or strong activation of OTX2 and absence of *GBX2::tdTOM* (yellow), **(k)** Hexbin density plots of per-cell mean intensity for *GBX2::tdTOM* and OTX2, median-normalised to the non-targeting reference condition and shown on a pseudo-log scale [log_10_(x+0.1)]. Solid/dashed lines = lower/upper classification thresholds (control-condition percentiles, per replicate); orange line = linear-regression fit (visual guide only). One representative replicate shown. More details in Methods. **(I)** Spearman’s rank correlation (r) between *GBX2::tdTOM* and OTX2 per-cell intensities, computed per biological replicate (points, shapes = replicates); bar and whiskers = mean ± s.e.m. Significance: two-tailed one-sample t-test of r_s vs 0. *P < 0.05, **P < 0.01, ***P < 0.001; ns, not significant. ICC quantification data in (b’, e’-h’, j’) are presented as mean ± s.e.m. Each point represents a biological replicate. Two-tailed unpaired t-tests were used with Welch’s correction for unequal variances. *P < 0.05, **P < 0.01, ***P < 0.001; ns, not significant (P ≥ 0.05). Scale bar - 50*μ*m.

A low WNT or RA signaling environment is also associated with anterior hindbrain fates; however, inhibition of these pathways (using the tankyrase inhibitor XAV939 and pan RAR inhibitor AGN193109, respectively) did not result in expression of EN1 (Fig. 5e’), although there were signs of anteriorization to r2/r3-like lineages in “bump” cultures (see below). Treatment of “no bump” cultures with 3.0µM CHIR from D1-4 resulted in a mixture of midbrain-like (OTX2+/EN1+) and r1-like (*GBX2::tdTOM*+/EN1+) identities (Fig. 5e), further demonstrating that the window for posterior transition has not yet closed in D1 “no bump” cultures. However, delaying this treatment to D2-4 resulted in most cells retaining an OTX2 positive state (Fig. 5f). This pattern was also seen in Fig. 3d when testing the vagal competence window. Treatment of “bump” cultures with 3.0µM CHIR from D1-3 resulted in transition to spinal cord-like identity (Fig. 5g), which was not a surprise given this treatment is similar to our “long bump” condition (Fig. 3-4).

The genes *EGR2* (*Krox20*) and *MAFB* (*Kreisler*) exhibit a distinct pattern of expression across the hindbrain such that between them they can be used to positively identify rhombomere 3-like (r3; EGR+MAFB-), rhombomere 5-like (r5;EGR2+MAFB+) and rhombomere 6-like (r6; EGR2-MAFB+) lineages (Fig. 5c)^111,113,128–130^. In addition, *HOXB1* is expressed in rhombomere 4 (r4) and regions posterior to this position. This means that in conjunction with EN1 and GBX2 it is possible to putatively identify r1-6-like identities in culture (with rhombomere 2 identified as being negative for all named markers; Fig. 5a). Using this approach, we found that in the WNT “bump” condition, when followed by DSi only, cultures consisted of a mix of r2 and r3 lineages (Fig. 5g-h). When WNT “bump” was followed by inhibition of WNT signaling from D1-3 we observed a significant reduction of r3-like EGR2+ cells in favor of r2-like identity (Fig. 5h). RA-receptor inhibition did not appear to change the proportion of spatial lineages when compared to DSi only, but FGF8 treatment resulted in a shift in favor of HOXB1+ r4 cells (25%; Fig. 5g-h). The 0.9µM CHIR treatment condition resulted in a significant increase in HOXB1+ r4 cells (∼60%) representing a more powerful posteriorization effect than FGF8. MAFB was only seen in a small proportion (∼1%) of these (Fig. 5h). However, given the apparent posteriorizing effect of WNT activation between D1-3 within the hindbrain domain, our expectation is that higher concentrations (∼1.0-2.5µM) would elicit more MAFB positive cells. Despite the low MAFB numbers, both r5 and r6 cells could be identified (Fig. 5h). None of these markers were seen expressed in cultures derived from the “no bump” condition (Fig. 5e-h).

We next sought to determine if *GBX2* induction alone (without WNT “bump”) is sufficient to drive the transition to posterior identity by using CRISPRa as described previously^131^ (in either DSi + 0.9µM CHIR or NC induction conditions; Fig. 5i-l, Supp. Fig 9). Activation of *GBX2* expression at the start of differentiation increased the proportion of *GBX2::tdTOM* positive cells from about 20% to 50% in the “no bump” condition (at D2; Fig. 5j), but did not significantly decrease OTX2 positive cells. Instead, a large proportion of the culture exhibited coexpression of the two factors, although in cases of very high *GBX2::tdTOM*, OTX2 rarely co-localized (Fig. 5j-k). Targeting *OTX2* for activation in WNT “bump” cultures gave mirroring results with successful induction in some cells but not significant decrease in *GBX2::tdTOM* expression (Fig. 5j). To further understand the relationship between these two factors we plotted their normalized intensity and found a significant negative correlation between them (spearman r ∼-0.5; Fig. 5k-l). Together these results support the mutually repressive role proposed by others^96–99,122,124,125,132,133^ and supported by evidence presented in Figure 4. However, further differentiation under DSi conditions to D4 did not result in significant changes of rhombomere-specific markers EGR2, MAFB or HOXB1 (Supp. Fig. 9a-b) and repeating the experiment for NCC differentiation conditions did not result in a significant change in *HOXB5* or *PHOX2B* expression (Supp. Fig. 9c-e).

### Applying patterning insights to midbrain dopaminergic neuron differentiation

Midbrain dopaminergic (mDA) neuron differentiation is thought to require low levels of WNT activation within the first two days of hPSC differentiation^62–66^. The rationale for this approach is the posteriorization of cells away from forebrain fates and toward midbrain fates. However, using this strategy can sometimes result in the generation of hindbrain or even spinal cord derivatives in unpredictable amounts, as well as other off-target lineages ^70,71^. The presence of off-target cell types in these cultures is of particular concern in the generation of hPSC-derived mDA neurons for the treatment of Parkinson’s disease, a strategy that has recently shown promise in Phase 1 clinical trials ^7,8^. Our results suggest that a barrier forms blocking posterior transition at around 48hrs (Fig. 3d, 5g). WNT activation before this point has the potential to direct hindbrain or spinal cord fate, while WNT activation after this point should only posteriorize within the anterior OTX2+ cranial compartment. To test this, we adapted our previously described mDA differentiation protocol such that WNT signaling was inhibited for a period of 48-72hrs (using 2µM XAV939; “inverted bump”) before being activated as usual (1µM CHIR; Fig. 6a). Using this strategy on optimal cell lines (KOLF and H9), we found comparable mDA progenitor generation at D11 (OTX2+EN1+FOXA2+; Fig. 6b). We found that the proportion of PITX3 positive cells at D25 was equivalent to controls (Fig. 6c) and that the efficient generation of TH+ FOXA2+ PITX3+ dopaminergic neurons was comparable at D32 (Fig. 6d). Using the H9-*GBX2::tdTOM* reporter we identified a decrease in the amount of contaminating GBX2+ lineages that was proportional to the duration of XAV treatment (day 6; Fig. 6e). We next tested this “inverted bump” approach with a 6-cell line pool to represent the functional variation exhibited across different hPSCs ^134,135^. Using this 6-cell line pool we observed a proportion of CDX2+ cells in control cultures that were not present when using the “inverted bump” strategy (day 5; Fig. 6f). At day 11, bulk RNA sequencing showed a reduction in *GBX2* and *CDX2* expression, and an increase in *OTX2* expression, in the “inverted bump” condition (KOLF; Fig. 6g). Finally, when using the 6-cell line pool, day 37 cultures exhibited a higher proportion of neurons positive for FOXA2 and TH in the “inverted bump” condition (Fig. 6h). These results provide a platform for increased robustness of mDA neuron generation across hPSC lines and further confirm the primary axis patterning principles identified for differentiating hPSCs in this study.

**Figure 6.**
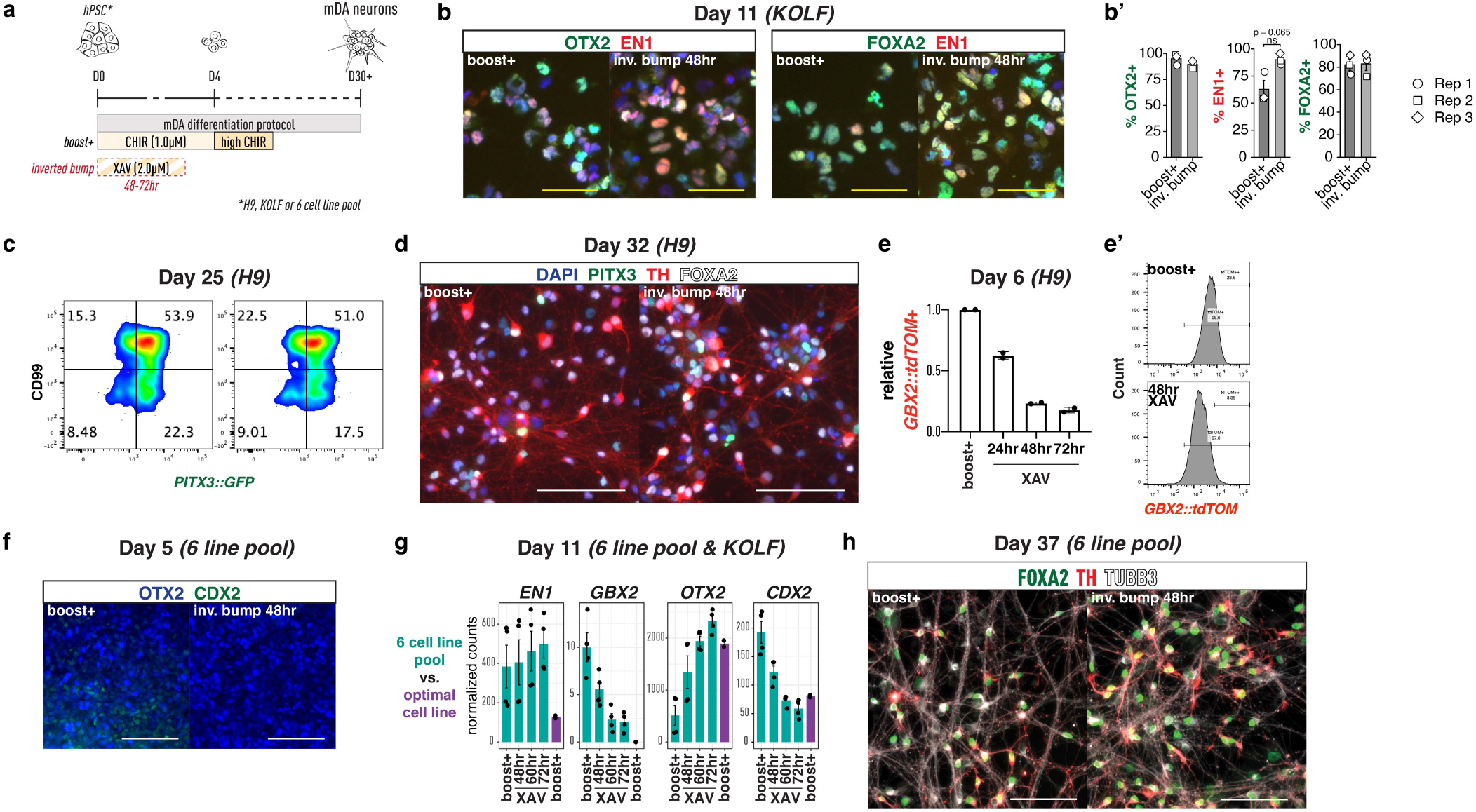
Inhibition of WNT signaling until the posterior competence window has closed results in a reduction of off target derivatives in midbrain dopaminergic neuron differentiation, **(a)** Schematic showing experimental “inverted bump” mDA protocol. Changes from previous “boost+” protocol (Kim et al., 2021; 2026) highlighted in red. **(b)** ICC at day 11 showing expression of mDA progenitor markers. Quantification in (b’). **(c)** Flow cytometry for CD99 and *PITX3::GFP* (using *H9-PITX3::GFP* reporter cell line), **(d)** ICC at day 30 showing expression of mDA neural markers *PITX3::GFP,* TH and FOXA2. **(e)** Summary flow cytometry data showing proportion of *GBX2::tdT0M+* off target cells at day 6 in different conditions. Representative plots in (e’). (f) Representative ICC showing off-target CDX2+ cells at day 5 of mDA differentiation with a 6 cell line pool, (g) Bulk RNA-seq data showing expression of axial markers at day 11 of mDA differentiation across different conditions in the 6 cell line pool and an optimal cell line (KOLF; only boost÷ shown), (h) Representative ICC for day 37 dopaminergic neurons expressing *F0XA2, TH* and *TUBB3.* Scale bar: White - WOpm; yellow - 50*μ*m.

### 3D assembly demonstrates “locked in” anterior-posterior identity

To test the degree of commitment of early progenitors to specific primary axis identities we conducted 3D co-culture experiments, aggregating progenitor cultures after 24hr (or 48hr for “long bump”) before assembling them at D2 into triple-origin assembloids and subjecting them to axial patterning factors alongside DSi (Fig. 7a). We did this using H9-derived hPSC cell lines with three distinct nuclear-localized fluorescent tags inserted into the locus of the housekeeping gene GPI, as has been done previously ^136^. H2B-GFP was used for anterior “no bump” cultures, H2B-tdTomato was used for posterior (hindbrain-like) WNT “bump” cultures, and H2B-3xFLAG was used for posterior (trunk-like) “long bump” cultures. We found that exposing these assembloids to high levels of CHIR (3µM) from D2-4 did not result in loss of *OTX2* expression in the “no bump”-derived lineage, but did result in gain of EN1 expression, indicating posteriorization, but only within the anterior cranial domain (Fig. 7b). Conversely, inhibition of WNT signaling with XAV (2µM) did not result in gain of *OTX2* expression in the WNT “bump”-derived lineage, confirming that 24hr WNT activation engages posterior fate irreversibly. High levels of CHIR resulted in acquisition of *CDX2* expression in the WNT “bump”-derived lineage, suggesting that at D2 these cells still have the potential to transition to trunk fates (Fig. 7b). Only treatment with RA appeared to break the anterior progenitor “lock in”, inducing a loss of *OTX2* and gain of *HOXB1* and *CDX2* expression in some of the “no bump” derived cells (Fig. 7b).

**Figure 7.**
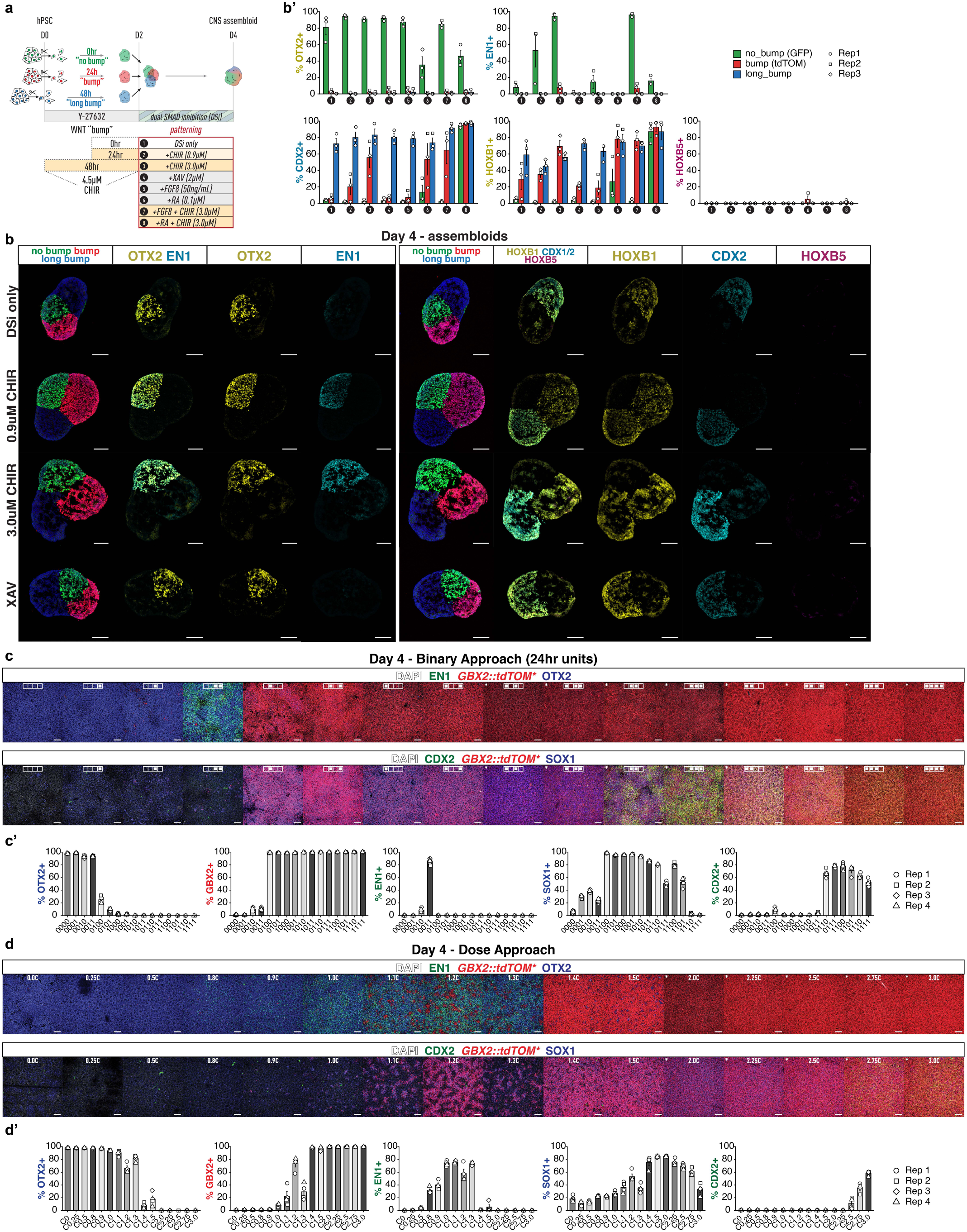
Anterior and posterior cranial progenitors show “locked in” regional commitment in a 3D assembloid system and a binary WNT-based approach to axial patterning generates all major regions, **(a)** Schematic showing experimental design of assembloid studies, **(b)** Representative ICC showing sections of day 4 assembloids. WNT-based patterning examples are presented (1 condition per row) with quantification by progenitor type for all conditions tested in (b’). The first column for each staining panel depicts the GFP/tdTOM stain that allows for identification of the progenitor type of origin, **(c-d)** Representative ICC from comparison of a binary axial patterning approach using maximum WNT activation (4.5pM CHIR) or WNT inhibition (2pM XAV) in 24hr blocks (c) with a traditional dose response approach titering CHIR concentration from 0-4.5/7M CHIR (d). In (c) the inset white symbol depicts the condition with a solid white square representing WNT activation and an empty square representing WNT inhibition. ICC quantification in (b’) are presented as mean ± s.e.m. Each point represents a biological replicate - the mean of a representative section of 4-6 individual assembloids functioning as technical replicates. The percentage of positive cells was calculated within each progenitor group: no_bump (GFP^+^), bump (tdTomato^+^) and long_bump (untagged/FLAG), shown as adjacent bars colour-coded green, red and blue, respectively.ICC quantification data in (c’, d’) are presented as mean ± s.e.m. Each point represents a technical replicate. Scale bar - 100μM.

### An integrated binary model of primary axis patterning

Our data indicate that by manipulating WNT signaling in a binary manner and taking advantage of the loss of competency in specific axial states at predictable times, we can generate neural ectoderm derivatives exhibiting characteristics of all major domains of the nervous system. To demonstrate that this logic enables the generation of all major AP positional identities in a single experiment, we tested all possible combinations of either 4.5µM CHIR or 2µM XAV treatment in 24hr units across a 4-day period of dual SMAD inhibition (Fig. 7c). Using this approach, we could reproducibly generate near pure cultures of forebrain-, midbrain-, hindbrain-, and spinal cord-like CNS lineages (Fig. 7c). In contrast, a traditional CHIR dose response treatment resulted in increased variability and generation of mixed cultures, particularly when generating lineages at the level of the midbrain and hindbrain regions. (Fig. 7d).

## Discussion

In 1958 Sulo Toivonen, in reference to his own experiments on the processes of neural induction and posteriorization, wrote *“mesodermalization* [posteriorization] *does not require preliminary activation* [neural induction]*“*^137^. Pieter Nieuwkoop, perhaps restricted by his own terminology, rejected this saying: *“It cannot, however, be concluded from* [Toivonen’s] *experiments that both actions actually took place simultaneously or even in a reversed order; the transforming* [posteriorizing] *principle could have been “stored” until the process of activation* [neural induction] *had occurred”*^138^. Despite evidence from Toivonen and others ^137,139,140^, the prevailing view for decades was that *“Early in differentiation, all neural cells have a rostral character”*^141^, which can later become reprogrammed posteriorly (although some questioned this perspective^142^). Work from the Briscoe lab^44^, building off extensive evidence from mouse development and human/mouse stem cell models ^2,6,29–46^, challenged this model by demonstrating in the mouse that trunk neural tissues emanate from a pre-neural posterior epiblast progenitor (NMP), supporting Toivonen’s thesis above. However, their study left hindbrain patterning embedded in the old default-anterior neural induction model.

Our data indicate that posterior cranial (hindbrain-level) identity is also established by a committed pre-neural progenitor, determined by specific WNT signaling dynamics alone, which functions as an intermediate step towards trunk identity (Fig. 8a). This sufficiency of WNT signaling is unsurprising: WNT’s role in primary axis patterning, complementary to that of the Hox genes, dates back to pre-bilaterians (Cnidarians)^18,143–146^. Orthologs for the vertebrate *OTX*, *GBX* and *CDX*^146–149^ genes also emerged at this point, as did BMP’s role in patterning the secondary (dorso-ventral) axis^150^. Other major pathways associated with primary axis patterning adopted this role later in evolution. For example, in Cnidarians asymmetrical expression of FGFs and FGF receptors is related to formation of the apical sensory organ (at the opposite pole to the WNT signaling center)^151,152^, while retinoic acid is not known to play a role in their development. The retinoic acid receptor (RAR) evolved later as a bilaterian innovation, initially functioning as a low affinity sensor for neural induction in protostomes before adopting an axial patterning role in chordates^18,153^.

**Figure 8.**
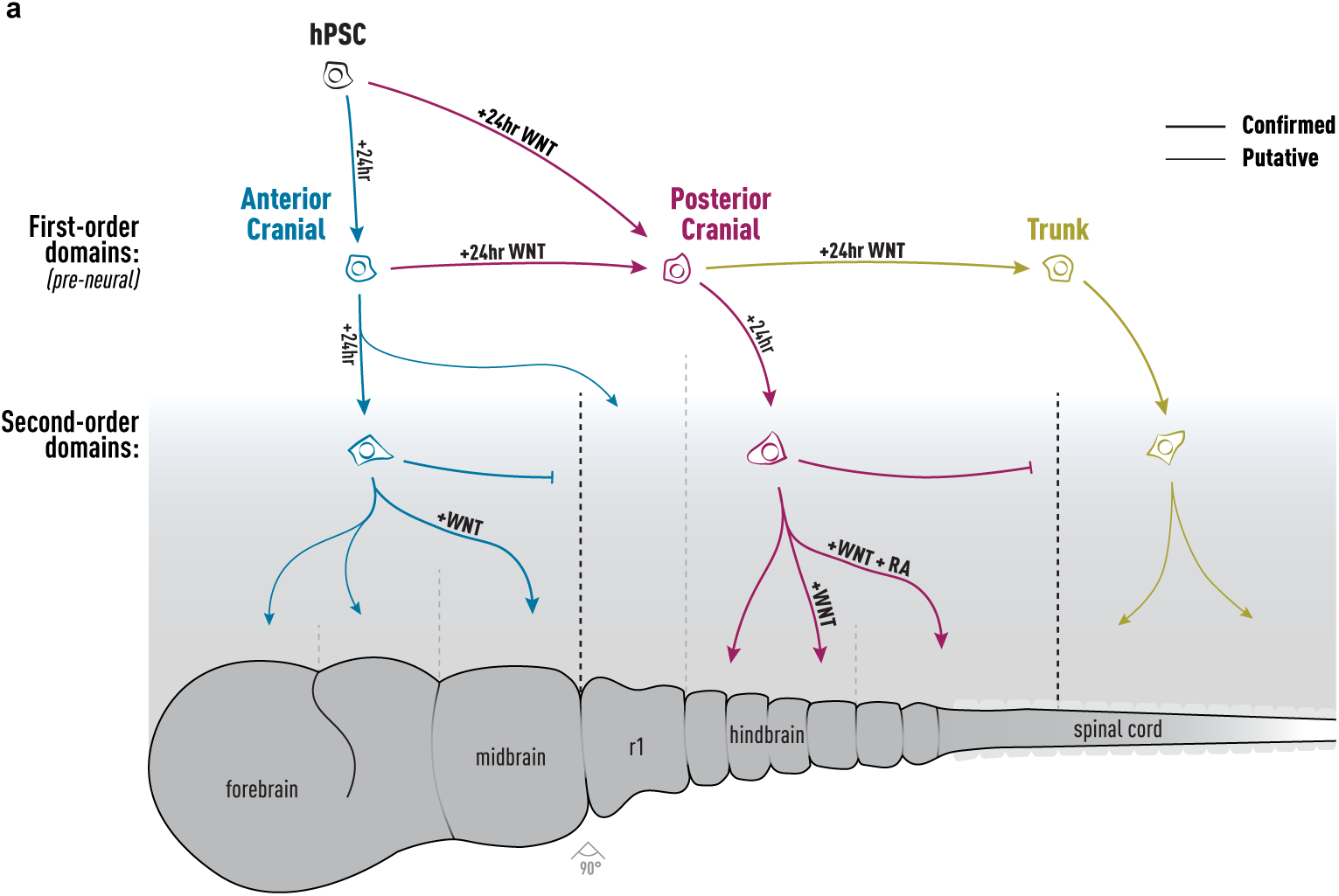
A new model of primary axis induction that describes a “third” axial progenitor type that exists as a transition state between anterior cranial and trunk identities but can also generate ectodermal derivatives in the hindbrain region. **(a)** Schematic representing fate decisions during early primary axis patterning in differentiating hPSCs. Axial fate decisions are organized hierarchically; first-order domains “Anterior Cranial”, “Posterior Cranial” and “Trunk” are specified through specific WNT dynamics and then second-order domains such as midbrain or distinct rhombomeres are specified through the continued action of WNT alongside FGFs and retinoic acid (RA). Fate barriers are established during differentiation that delineate competence windows within which WNT and other signals exhibit defined roles. Developmental pathways and fate barriers are presented with a thick line if confirmed in this study and a thin line (and grey in the case of barriers) if they are putative. r1, rhombomere 1; +WNT, represents treatment with CHIR99021; RA, retinoic acid.

With the understanding that distinct dynamics of WNT activation are sufficient for hindbrain-level lineages as well as trunk we can begin to describe an integrated model of patterning down the entire neural axis: after early commitment to first-order axial domains (“anterior cranial”, “posterior cranial” and “trunk”), second-order domains (e.g. forebrain vs. midbrain or distinct rhombomeres) can be subsequently specified by WNT and other factors (e.g. RA) in a nested fashion (Fig. 8a). This hierarchy resembles the broad precursor domains to the canonical 11 progenitor regions that emerge in dorso-ventral patterning in the spinal cord^154^. Posterior cranial progenitors from the “bump” condition generate all hindbrain levels except rhombomere 1-like (r1) lineages; we present specific conditions that enrich for r2-8-like lineages from these progenitors. r1 lineages were only seen with late WNT activation in “no bump” cultures before anterior commitment was fully established (Fig. 8a). This raises the intriguing possibility that r1 lineages are developmentally more closely related to anterior tissues than the rest of the hindbrain, although further work is needed to confirm this. The vagal region may also constitute a first-order domain, as 18hr WNT activation was sufficient to induce GBX2 expression but not vagal competence, which required 24hr. Recent work from the Metzis lab supports this, implicating *CDX1* expression as a requirement for vagal identity^93^. We observe *CDX1* transcripts in our scRNA-seq data at D1 but have yet to confirm its absence at 18hr. Recent work by the Loh lab has demonstrated that posterior hindbrain-level CNS derivatives can be generated from hPSCs via FGF and RA activation alone, without WNT^155^. This suggests that for certain lineages the posterior cranial commitment event we observed in our genetic fate mapping experiments can be bypassed, presumably through accessibility of lineage-specific coordinators to RA and FGF signal response machinery. Indeed, in our assembloid experiments we observed that the anterior cranial commitment established at 48-72hr can be overridden by RA treatment, resulting in the loss of OTX2 (Fig. 7b’). This effect is likely limited to the retinoid-dependent part of the hindbrain^20,129^. Furthermore, with the RA and FGF approach posterior progenitors appear to be CNS restricted and unable to transition to trunk fates, additional features of a later stage identity ^44,155^.

The transition to posterior fate is coordinated by WNT response elements alongside Homeobox (ANTP & POU) class transcription factors. Our data support a well described mechanism whereby TCF7L1/2 suppress WNT targets by association with the TCF/LEF consensus sequence. Upon canonical WNT activation, TCF7/LEF1 associate with β-catenin, displacing TCF7L1/2 and activating the posterior GRN ^102–107^. Functionally, once posterior fate is established (∼18hr WNT activation), the anterior cranial GRN cannot be re-activated by removing CHIR. This is likely due to epigenetic suppression of network members such as OTX2, as shown in our CUT&RUN analysis. Commitment to anterior cranial fate, which emerges after the period of posterior competence elapses, likely functions in a similar way where the putative poised state we observe for genes like *GBX2* at D1, shifts to inactive after 48-60hr of WNT inhibition.

The antagonistic regulatory relationship between OTX2 and GBX2 in coordinating axial fate has been reported on extensively^96–99,122,124,125,132,133^. Our study supports this, with inverse expression patterns, mutual binding of loci, and transient repression of each other after CRISPRa activation. However, early transient OTX2 activation did not block vagal competence or hindbrain fate, nor did GBX2 activation significantly impact anterior identity. In the mouse, *Otx2* is known to be regulated by a suite of stage-specific enhancers^97,156,157^. Much attention has been given to the anterior neural (AN) and forebrain/midbrain enhancers (FM and FM2), where Gbx2 restricts *Otx2* expression through competition with Pou factors (such as Pou3f1, the human orthologue of which was identified in our SCENIC analysis)^97,156^. However, these enhancers activate at E7.0 (AN) and E8.0 (FM/FM2), but anterior restriction of *Otx2* begins at E6.5^157–159^. *Gbx2* is not yet expressed at this stage, but the epiblast (EP) *Otx2* enhancer (part of the broader region around AN) is expressed, and transitions from marking the whole epiblast to anteriorly restricted activity^157^. This suggests that the first step of *Otx2* repression might be Gbx2-independent, consistent with GBX2 knock outs losing r1-3, but not the whole hindbrain^122,160–165^.

*GBX2*, alongside *CDX1* and *CDX2*, are known to be direct targets of WNT signaling, so their upregulation at D1 in the “bump” condition is expected^87,166,167^. Despite early detectable *CDX2* transcripts, by day 2 any indication of trunk identity is lost from “bump” cultures, sustained only where WNT activation has been maintained (“long bump” cultures). This is reminiscent of the culture-wide *GBX2::tdTOM* induction seen after 12 hours of WNT activation, while 18hr or more is required for *GBX2::tdTOM* activity to be fully established and not lost by D4. The capacity of the “bump” progenitors to generate hindbrain and vagal lineages supports their non-trunk identity, but the expression of TBXT and CDX2 at D2 in “long bump” conditions indicates retained trunk potential. In the extensive literature about NMP specification, little focus has been given to the possibility that NMPs may be derived from a pre-neural posterior cranial progenitor. However, expression of GBX1/2 extends into the spinal cord and GBX2 lineage tracing experiments identify the spinal cord as being GBX2-derived, supporting this model of _development_122,155,160–165.

Temporally restricted, saturating WNT activation can generate the same axial lineages as concentration-dependent dosing, suggesting the WNT integral within a restricted timeframe is key for fate specification. Our binary approach seems particularly useful in border zones, where dosing methods often result in mixed identity cultures Robust, high-efficiency protocols are essential for translating hPSC technology into clinically successful regenerative therapies. Recent mDA neuron transplantation trials for Parkinson’s Disease (PD) have shown a good safety profile and early signs of efficacy ^7–9,168^. However, some early efforts to treat PD with fetal-derived mDA neuron progenitors resulted in dyskinesia caused by contaminating serotonergic neurons, possibly carried in the graft from the donor fetal hindbrain^169,170^. Benchmarking work integrating scRNA-seq data from current mDA stem cell models identified off-target cell types, including V3/floor plate hindbrain lineages, though minimal serotonergic contamination ^71^. This demonstrates the need for more robust axial patterning in mDA differentiation systems to avoid off-target cell types and to facilitate autologous approaches where in-depth testing is more challenging compared to allogeneic, off-the-shelf approaches.

The other differentiation system targeted for improvement in this project, hPSC-derived enteric neural crest, presents a promising cell replacement therapy candidate for enteric neuropathies, including Hirschsprung’s Disease (HD), esophageal achalasia and gastroparesis^171^. Current surgical treatment for HD is lifesaving, but causes major lifelong complications with 95% readmission rates^172^. Transplanting hPSC-derived enteric NCCs into HD mouse models and human gut tissue has shown extensive integration and functional restoration ^4–6,11^. However, as with the mDA neuron progenitors, off-target lineage contamination remains a barrier. Contaminating anterior neural crest progenitors can yield mesenchymal derivatives (bone, cartilage, smooth muscle) that may cause unexpected side effects.

Finally, because second-order domains within the hindbrain are defined by discrete rhombomeric identities, a patterning logic that resolves the axis at this granularity offers a rational entry point to lineages that remain difficult to access. Future studies may enable the selective derivation of branchiomotor and visceromotor neurons (r2-r8)^173^ and of rostral versus caudal serotonergic raphe neurons (r1-3, r5-7)^174^, or the more selective generation of cerebellar lineages (r1)^175,176^. Similarly, NCCs exhibit a high degree of fate restriction by axial identity^177^, and not all subtypes can yet be generated from hPSCs despite recent advances^74^. Our work could facilitate efforts to derive cardiac NCCs, which emerge adjacent to the vagal region targeted here and hold broad potential for disease modeling and cell therapy. More generally, the implementation of binary, time-based patterning approaches offers a general strategy for the efficient production of defined neural derivatives along the whole-body axis, and a route toward generating the complete atlas of human cell types on demand.

## Methods

### hPSC lines and cell culture

Experiments with hPSCs and iPSCs were in compliance with the Tri-Institutional ESCRO at Memorial Sloan Kettering Cancer Center, Rockefeller University and Weill Cornell Medicine. hESC lines WA09 (H9; 46XX) and WA01 (H1; 46XY) were from WiCell Stem cell Bank. Reporter lines H9-SOX10::GFP and H9-GBX2::tdTOM were derived as described previously ^90^. hiPSC line KOLF2.1J (KOLF; 46XY) is a well-characterized reference cell line ^178,179^ and the 6-cell line pool consists of healthy controls from the Parkinsons Progression Marker Initiative (PPMI)^134,135^. All hiPSCs were grown in E8 medium (Gibco - A1517001) on Laminin-521 (Gibco - 29248) or vitronectin (mDA neuron experiments; Thermo Fisher, A14700) and passaged every 3-4 days using EDTA (0.5 M EDTA/PBS) and routinely tested for mycoplasma contamination.

### Generation of reporter lines

All hPSC reporter lines were generated by Cas9-mediated homology-directed repair (HDR) in H9 human embryonic stem cells as described previously^90^. The H9-SOX10::GFP reporter line was generated by in-frame knock-in of a P2A-H2B-GFP cassette into the SOX10 locus, as previously described^180^. To generate the H9-GBX2::tdTomato reporter line, an sgRNA targeting a sequence proximal to the stop codon of the GBX2 gene was designed and cloned into the pX330-U6-Chimeric_BB-CBh-hSpCas9 vector (Addgene plasmid #42230). A donor plasmid was constructed containing 400-bp left and right homology arms flanking a P2A-H2B-tdTomato cassette and a floxed neomycin-resistance cassette. The sgRNA/Cas9 plasmid and donor plasmid were co-electroporated into H9 cells using a Lonza 4D-Nucleofector with P3 Primary Cell Solution and pulse code CB-150. Neomycin (G418) selection was subsequently applied to enrich for targeted cells. The neomycin-resistance cassette was removed by Cre-mediated recombination by electroporation of 1 µg of a Cre-expressing plasmid into the cell population. Individual clones were subsequently isolated and screened by PCR to identify clones with the desired knock-in and successful Cre-mediated excision, which were further confirmed by Sanger sequencing (Supp. Fig. 10).

### Neural crest differentiation (“bump” protocol)

Neural crest differentiation (“bump” protocol) was carried out by dissociating hPSC cultures 3-4 days post-passage using Accutase (5 minutes at 37°C) and seeding cells down onto matrigel at 30,000 cells/cm^2^ in E6 medium supplemented with the ROCK inhibitor Y-27632 (10µM) and CHIR99021 (4.5µM). Exactly 24 hours later (D1) they were switched to neural crest induction medium consisting of DMEM F12, 1x N2 supplement, SB431542 (2µM), CHIR99021 (0.9µM), DMH1 (1.0µM) and BMP4 (15ng/mL). This medium uses the top-down inhibition strategy for controlling BMP activity developed previously ^181,182^. At day 4 (D4; when cultures reach ∼100% confluency) this medium was supplemented with 0.1µM RA. At day 7 (D7) cultures were ready for downstream analysis or differentiation. All cultures had media exchanged daily. Variations on the above including modulations to “bump” treatment, and supplementing the media with EDN3 (100nM) or GDNF (25ng/mL) from D4-7, are described in figure schematics and the text. For the CellTag experiment (Fig. 1) a low-density starting seed (20,000 cells/cm^2^) was used to moderate end point cell numbers and ensure representation of clones. Cultures were differentiated until day 9 (D9) to ensure expansion of vagal clones post-RA treatment. A stepwise protocol is available on request.

### ENS differentiation

ENS differentiation was carried out using the ENS medium described previously^5^. D7 vagal NCC were dissociated using Accutase (5 minutes at 37°C) and seeded onto Laminin/Fibronectin coated plates at a density of 80,000 cells/cm^2^ (“bump”) or 150,000 cells/cm^2^ (“no bump”) in ENS medium consisting of Neurobasal medium, 1x N2 supplement, 1x B27 supplement (with Vitamin A), GDNF (25ng/mL), 1x Glutamax, ascorbic acid (100µM) and Y-27632 (10µM). After 24 hours Y-27632 was removed. Media was changed daily until day 40 (D40) when cells were taken for analysis. Ensure high media volume between days 14-28 (D14-28) when “bump”-derived ENS precursors are highly proliferative and metabolically active. D40 cultures were dissociated using Accutase supplemented with papain containing neural isolation enzyme at 1:50^183^(Neuron Isolation Enzyme for Pierce; Thermo Scientific, 88285). For this, gentle trituration was used with a P1000 every 30 minutes until clumps were not visible by eye. At each 30-minute interval the supernatant was taken and neutralized in ENS medium on ice while the dissociation was continued with fresh Accutase papain mix. A stepwise protocol is available on request.

### Dual SMAD inhibition differentiation

The dual SMAD inhibition experiments presented in Figure 5 were carried out by dissociating hPSC cultures 3-4 days post-passage using Accutase (5 minutes at 37°C) and seeding cells down onto matrigel at 30,000 cells/cm^2^ in E6 medium supplemented with the ROCK inhibitor Y-27632 (10µM). These cultures were subjected to variations of the “bump” treatment (4.5µM CHIR99021 for 0-24 hours) as described in the schematic and the text. Exactly 24 hours later (D1) the media was exchanged for DSi medium consisting of Neurobasal, 1x N2 supplement, 1x B27 supplement (without Vitamin A), 1x Glutamax, LDN193189 (100nM; Stemgent) and SB431542 (10µM). Cells were maintained in these conditions with further patterning agents as described in the schematic and text with media changed daily until D4, when they were analyzed.

### mDA neuron differentiation (“inverted bump” protocol)

Midbrain dopaminergic (mDA) neuron differentiation was carried out according to an adapted version of the Boost+ protocol^62,63^ including the “inverted bump” modification described in Figure 6. Briefly, hPSCs were dissociated using Accutase (5 minutes at 37°C) and seeded onto Geltrex (1:40)-coated plates at 500,000 cells/cm² in day 0 medium consisting of Neurobasal, 1x N2 supplement, 1x B27 supplement, Glutamax, Pen-Strep, LDN193189 (250nM), SB431542 (10µM), SAG (2µM), XAV939 (2µM) and Y-27632 (10µM). Cultures reached 100% confluency by day D1, at which point Y-27632 was withdrawn. Cells were fed daily with the same medium on D2. XAV was withdrawn after 48-72 hours as described in the figure and the text. At the point of XAV withdrawal CHIR99021 (1µM) was added. At D4, CHIR99021 was boosted to 6µM (“CHIR boost”) while LDN193189, SB431542 and SAG were maintained, and cultures were fed daily with this medium through D6. At D7, LDN193189, SB431542 and SAG were withdrawn while the CHIR boost (6µM) was continued through D9. At D10, CHIR99021 was reduced to 3µM and cultures were fed with Neurobasal, 1x N2 supplement, 1x B27 supplement, Glutamax, Pen-Strep, BDNF (20ng/mL), ascorbic acid (0.2mM), GDNF (20ng/mL), dcAMP (0.5mM) and TGF-b3 (1ng/mL). At D11, cultures were dissociated using Accutase (30–40 minutes at 37°C), filtered through a 40um strainer, and replated onto fibronectin and laminin-coated plates at 900,000 cells/cm² in D10 medium (3µM CHIR99021). From D12-15, cultures were matured in Neurobasal, 1x N2 supplement, 1x B27 supplement Glutamax, Pen-Strep, BDNF (20ng/mL), ascorbic acid (0.2mM), GDNF (20ng/mL), dcAMP (0.5mM), TGF-b3 (1ng/mL), IWP2 (1µM) and FGF18 (100ng/mL). Cultures were passaged again at D16 by Accutase dissociation and replated onto fibronectin and laminin-coated plates at 900,000 cells/cm², then maintained in the same maturation medium with DAPT (10µM) substituted for IWP2/FGF18 through D25. At D25, cultures underwent a final low-density replating step (Accutase dissociation, filtration, 200,000 cells/cm²) and were subsequently maintained in maturation medium with feeds every 3 days. A stepwise protocol is available on request.

### Assembloid experiment

The assembloid experiment was carried out by dissociating hPSC cultures 3-4 days post-passage using Accutase (5 minutes at 37°C) and seeding cells down onto matrigel at 30,000 cells/cm^2^ in E6 medium supplemented with the ROCK inhibitor Y-27632 (10µM) and with or without CHIR99021 (4.5µM) according to the condition. The “long bump” condition was started one day earlier (D-1) so that the second day of CHIR treatment would still occur in monolayer format to keep it consistent with previous experiments. At D1 aggregation was carried out by dissociating cultures with Accutase (5 minutes at 37°C) and seeded into non-adherent v-bottom 96-well plates (PrimeSurface; Sbio, MS-9096VZ) at 2000 cells/well for “bump” and “long bump” and 3000 cells/well for “no bump” (due to reduced survival in the absence of CHIR) in E6 medium supplemented with the ROCK inhibitor Y-27632 (10µM). 50uL of media was used per well and 1 plate per condition. At D2 assembly was carried out by transfer of aggregates into a single plate row-by-row with a multichannel pipette. Aggregates were left to assemble for 2 hours before the media was exchanged for patterning medium, E6 supplemented according to the schematic in Figure 7. This medium was refreshed on D3. On D4 assembloids were taken for analysis.

### Binary axial patterning approach

The integrated binary axial patterning system presented in Figure 7 was carried out by dissociating hPSC cultures 3-4 days post-passage using Accutase (5 minutes at 37°C) and seeding cells down in 384-well plates onto matrigel at 300,000 cells/cm^2^ in E6 medium supplemented with the ROCK inhibitor Y-27632 (10µM) with or without CHIR99021 according to the condition. Cultures were fed daily and analyzed at D4. A stepwise protocol for specific regions is available on request.

### CRISPRa experiment

Activation of endogenous *GBX2* or *OTX2* was performed as previously described^131,184^. Briefly, sgRNAs targeting *GBX2* (TGGCCCGGCTCGCCTCACGT) or *OTX2* (ATTGACACATCTAAGCCAGA) were designed. An sgRNA targeting CTGAAAAAGGAAGGAGTTGA was used as a non-targeting control. SAM-TET1 components together with the corresponding sgRNA plasmid were introduced into cells by electroporation, along with a puromycin-resistance plasmid for enrichment of transfected cells. Cells were selected with 0.5 µg/mL puromycin in culture medium beginning 24 h after electroporation for 1 day.

### Immunocytochemistry

Cultures were fixed for 10 minutes in 4% PFA PBS. They were washed twice in PBS and then permeabilized in PBST (PBS 0.5% Triton X-100). They were then blocked for 1 hour in Blocking Buffer (PBST 5% donkey serum) before undergoing primary antibody stain overnight in blocking buffer at the concentrations listed in Table 1. The following day they were washed 3x in PBST over the course of 1 hour and then incubated in secondary antibody solution for 1 hour. They were then subjected to a further 3 washes in PBST over the course of an hour and a 30-minute incubation with DAPI before imaging using a Zeiss Axio Observer microscope.

**Table 1.**

| Target | Species/Isotype | Dilution | Source/Catalog # | Clone | Conjugate | Marker of.. | Application |
| --- | --- | --- | --- | --- | --- | --- | --- |
| anti-FLAG | ms/IgG1 | 1:500 | millipore/F1804 | M2 |  | n/a | ICC |
| Bra/T/TBX-T | GT | 1:200 | R&D/AF2085 | poly |  | Trunk/NMP | ICC |
| CDX2 | ms/IgG1 | 1:25 | DSHB/PCRP-CDX2-1A3 | 1A3 |  | Trunk/NMP | ICC |
| CGRP | ms/IgG2a | 1:1000 | abcam/ab81887 | 4901 |  | ENS | ICC |
| CHAT | Gt | 1:100 | millipore/AB144P | poly |  | ENS | ICC |
| EGR2 | ms/IgG1 | 1:100 | sc-518117 | G-9 |  | Hindbrain | ICC |
| EN1 | Rb | 1:300 | novus/NBP2-57757 | poly |  | mDA, Midbrain, R1_Hindbrain | ICC |
| FOXA2 | GT | 1:200 | R&D/AF2400 | poly |  | mDA | ICC |
| GBX2 | RB | 1:25 (C&R) | proteintech/21639 | poly |  | Hindbrain | CUT&RUN |
| GFP | CHK | 1:2000 | ab13970 | poly |  | n/a | ICC |
| HOXB1 | SHP | 1:200 | R&D/AF6318 | poly |  | Hindbrain | ICC |
| HOXB4 | rat/IgG2a | 1:25 | DSHB | I12 |  | Hindbrain | ICC |
| HOXB5 | Rb | 1:200 | ab254882 | poly |  | Hindbrain | ICC |
| MAFB | Rb | 1:500 | HPA005653 | poly |  | Hindbrain | ICC |
| Neuromedin U (NMU) | Rb | 1:200 | St John's Laboratory Ltd/STJ116312 | poly |  | ENS | ICC |
| NOS1 (nNOS) | Rb | 1:500 | millipore-07-571 | poly |  | ENS | ICC |
| OTX2 | GT | 1:500(ICC)<br>1:25(C&R) | R&D/AF1979 | poly |  | Forebrain/Midbrain | ICC, CUT&RUN |
| p75 | ms/IgG1 | 1:200 | biolgd/345107 | ME20.4 | APC | Neural Crest | Flow Cytometry |
| PAX6 | msIgG2a | 1:200 | BD pharmingen/561462 | O18-1330 |  | CNS | ICC |
| PENK | ms/IgG1 | 1:300 | abcam/ab150346 | NOC1 |  | ENS | ICC |
| RFP | ms/IgG2a | 1:500 | Rockland/200-301-379 | 8E5.G7 |  | n/a | ICC |
| SSEA3 | rat/IgM | 1:200 | BD pharmingen/561145 | MC-631 | 647 | hPSC | Flow Cytometry |
| tdTomato | Rb | 1:400 | thermo/TA150089 | poly |  | n/a | ICC |
| TFAP2A | ms/IgG2b | 1:100 | sc-12726 | 3B5 |  | Neural Crest, NNE | ICC |
| TUBB3 | ms/IgG2a | 1:500 | biolgd/801201 | TUJ-1 |  | neurons | ICC |
| Tyrosine Hydrolase (TH) | ms/IgG1 | 1:300 | millipore/MAB318 | LNC1 |  | mDA | ICC |
| VIP | Guinea Pig | 1:200 | synsys/443005 | poly |  | ENS | ICC |
| H3K4me3 | Rb | 1:50 | abcam/ab8580 | poly |  | Active Transcription | CUT&RUN |
| H3K27me3 | Rb | 1:50 | Epicpypher/13-0055 | 2084-1G5 |  | Inactive Transcription | CUT&RUN |

Assembloids were fixed in 4% PFA PBS for 3 hours and then washed 3 times in PBS before cryoprotection in 15% sucrose overnight followed by 20% sucrose for a second night. They were then mounted in OCT and cryosectioned at 12µm thickness. After drying, sections were washed overnight in PBST. They were then subjected to overnight primary and overnight secondary antibody incubation with washing in PBST during the day. Assembloids were imaged using a Leica Stellaris confocal capable of imaging at 750nm enabling use of a 6^th^ channel. To adequately capture the whole section z-stacks of 3-5 images at 2um intervals were taken and a max intensity projection used for presentation.

### Image analysis and quantification

For all image analysis, segmentation of the DAPI channel was carried out using Ilastik. For standard cell culture ICC quantification, 9 approximately equally spaced fields of view were imaged per well for a minimum of 3 biological replicates. A custom FIJI macro was used to calculate the proportion of positive cells per field/channel/condition/replicate based on mean intensity values and thresholding set using negative controls. A custom R pipeline was used to plot data as bar charts (with a single point representing a biological replicate) and perform T tests. In the case of the CRISPRa experiments a separate FIJI macro was designed to record mean intensity per nucleus and the data plotted as median normalized intensity (each cell divided, per replicate, by the median of the non-targeting reference condition) on a scatter plot. Lower thresholds were calculated to lie at the 95^th^ percentile for OTX2 (calculated in the non-targeting “bump” condition where cultures are OTX2 negative) and the 75^th^ percentile for GBX2::tdTOM (calculated in the non-targeting “no bump” condition where approximately 20% of cells were tdTOM+). Upper thresholds were calculated on the basis of the upper 97.5^th^ percentile in the non-targeting conditions where the relevant marker is expressed. The line presented on scatter plot is a linear regression fit. Spearman’s rank correlation coefficient (r) was calculated independently of this line, per replicate. For assembloid image quantification a custom FIJI macro was used that first identifies progenitor origin according to GFP (“no bump”), tdTOM (“bump) or double negative (“long bump”) nuclei and markers were presented within each progenitor class. Due to some GFP/tdTOM weak cells getting mislabeled as “long bump” a local majority vote approach was used to correct within the “footprint” of the progenitor group.

### Flow cytometry

Neural crest differentiation efficiency was assessed by flow cytometry at day 9, using both high p75 reactivity and SOX10-GFP reporter expression to define the neural crest population. GBX2-tdTomato induction in response to canonical WNT activation was assessed by flow cytometry across days 0–1 of differentiation of the GBX2 reporter line, with SSEA3 used to assess pluripotency exit over this window. For both, cultures were dissociated using Accutase (5 minutes at 37°C) and neutralized in 3x volume of FACS buffer (PBS, 2% FBS). Cell suspensions were incubated for 30 minutes with anti-p75 or anti-SSEA3 antibody (see Table 1) and DAPI as a viability dye, then washed twice in FACS buffer before analysis on a BD LSRFortessa flow cytometer (BD Biosciences). Data were acquired using BD FACSDiva software (v9.0) and analyzed using FlowJo software (v11.1.1). Cells were gated on FSC/SSC to exclude debris, followed by doublet exclusion and DAPI-negative gating to select viable cells; positive/negative boundaries for p75, SOX10-GFP and SSEA3 were set using an unstained control.

### CellTag promoter cloning and lentiviral production

#### Library construction and promoter cloning

pSMAL (Addgene, 161785) was cut with Esp3I and NheI (NEB, R0734S and R3131S) to remove minimal CMV promoter. EF-1a Core Promoter was PCR amplified from LentiCRISPRv2 (Addgene, 52961), and restriction cloned into gel extracted Esp3I/NheI cut pSMAL. pSMAL-Ef1a Core Promoter plasmid was confirmed by sanger sequencing. Celltag V1, V2 and V3 barcodes (Addgene 115643,115644,115645) were then recloned into modified pSMAL-EF1a and barcode whitelist generated by NGS.

#### Virus production

HEK293T cells were maintained in DMEM supplemented with 10% FBS, 1× non-essential amino acids (NEAA), and 1× penicillin/streptomycin at 37 °C in 5% CO₂. One day before transfection, cells were reseeded in antibiotic-free medium (DMEM, 10% FBS, 1× NEAA) at 8-10×10⁶ cells per 10cm dish to reach 90-95% confluency. Cells were transfected with X-tremeGENE HP (Roche, 6366546001) using, per dish, 5 µg transfer plasmid (construct or library), 7.5 µg psPAX2, and 4.1 µg pMD2.G in Opti-MEM (Thermo Fisher, 31985062). Plasmid complexes were incubated for 15min at room temperature and added dropwise to cells in fresh antibiotic-free medium. Within 18h, medium was replaced with 7-8mL fresh medium. Supernatants were harvested at 48 and 72h post-transfection, pooled, clarified (1,250 × g, 10 min), and filtered (0.45 µm, PES). Clarified supernatant was combined with Lenti-X Concentrator (TaKaRa Bio, 631232) at 3:1 (supernatant:concentrator) and processed per the manufacturer’s instructions (1,500 × g, 45 min, 4 °C). Concentrated virus was aliquoted, snap-frozen, and stored at -80 °C.

### scRNA-seq sample processing and analysis

#### Sample prep

Library prep was carried out by the IGO or SAIL core facilities at MSKCC. Briefly, single cell suspension was loaded onto a Chromium Next GEM Chip G (10X Genomics #1000120) and GEM generation, cDNA synthesis, cDNA amplification, and library preparation of ∼10,000 cells per sample was carried out using the Chromium Next GEM Single Cell 3’ Kit v3.1 (10X Genomics #1000268) according to the manufacturer’s protocol. cDNA amplification included 11–12 cycles and 30–240 ng of the material was used to prepare sequencing libraries with 10–16 cycles of PCR. Indexed libraries were pooled equimolar and sequenced on a NovaSeq 6000 in a PE28/88 run using the NovaSeq 6000 S1 or S4 Reagent Kit (100 or 200 cycles) (Illumina). For the D4 and D7 scRNA-seq presented in Figure 2 antibody-based multiplexing was employed, using the antibodies in Table 1.

#### Data processing

Raw sequencing data were processed using Cell Ranger (10x Genomics) to generate feature-barcode count matrices. Downstream analysis was performed in R using the standard Seurat pipeline, including quality-control filtering of low-quality cells, log-normalization, identification of highly variable features, scaling, principal component analysis, graph-based clustering and UMAP-based dimensionality reduction.

#### Clonal labelling and clone calling

Cells were labelled with CellTag lentiviral libraries and clones were called essentially as described^77,185^, using CellTagR and the accompanying workflow (github.com/morris-lab/CellTagR). Reads were aligned with Cell Ranger to a custom human reference containing GFP and the CellTag UTR, and CellTag reads were recovered from the resulting BAM files. The CellTagV1 and CellTagV2 libraries were processed independently. CellTag UMI count matrices were error-corrected with Starcode, binarised at a threshold of ≥2 UMI per CellTag per cell, filtered against the corresponding library whitelist, and restricted to cells carrying between 3 and 20 CellTags. Clones were then called from the pairwise Jaccard similarity of cellular CellTag signatures using a correlation cutoff of 0.7, and clone identities were added to the Seurat object metadata. Of the 20,504 cells in the final dataset, 9,302 carried a CellTagV1 clone and 3,432 a CellTagV2 clone.

#### Clonal coupling analysis

Clonal relationships between transcriptional clusters were quantified as clonal coupling scores, following approach used by the Klein lab as implemented in Boezio et al. (2025)^154,186,187^. CellTagV1 and CellTagV2 clone assignments were analysed separately. Clone identifiers were made unique within replicate, only multicellular clones (≥2 cells) were retained, and clusters with fewer than 10 tagged cells were excluded. For each pair of clusters, observed clonal co-occurrence was calculated as the cross-product of the clone by cluster cell-count matrix, the diagonal of which gives the within-cluster term, the extent to which the cells of a clone remain within a single cluster. A null distribution was generated from 2,000 permutations in which cells were randomly reassigned to clones while preserving the number of clones, the clone size distribution and the number of cells in each cluster. Coupling scores are reported as z-scores of the observed co-occurrence relative to this null (positive, clonally coupled; negative, clonally segregated) together with two-sided empirical P values. Because clusters are largely timepoint-specific, cluster labels were permuted within timepoint, so that coupling between D4 and D9 clusters is evaluated against a null in which a D4 cell cannot be assigned a D9 identity. Heatmap rows and columns were ordered by hierarchical clustering on the correlation distance between coupling-score profiles using average linkage^186^, with the diagonal excluded from the distance calculation. Colour scales were set from the off-diagonal range of each matrix and values clipped to it. Directed D4 to D9 coupling was computed in the same way, using only clones with cells at both timepoints, with D4 clusters as rows and D9 clusters as columns and cluster labels permuted independently within each timepoint.

#### Cell type annotation with CellTypist

Cell type classification was performed using CellTypist (v2.0). Fetal and adult reference datasets were assembled from previously published single-cell RNA-sequencing studies ^80,81^ and annotated using their corresponding published cell type labels. Fetal and adult reference models were generated by training CellTypist classifiers with feature selection enabled to identify informative genes for model construction. For the adult reference, broad cell class labels were created by collapsing subtype-specific annotations into the major classes PIMN, PEMN, PSVN, PIN, PSN, and Glia. *In vitro*-derived cells were subsequently annotated using the corresponding pretrained fetal and adult CellTypist reference models. Cell identities were assigned using both best match, which assigns each cell to the highest-scoring reference cell type, and majority voting, which refines predictions based on the consensus annotation of neighboring transcriptionally similar cells.

### SCENIC analysis

The GRN inference package SCENIC^94^ was run using the pyscenic version^95^ with NES threshold = 2 for step 2. It should be noted that we initially found a large number of regulons were being dropped from the analysis during step 2 (Rcistarget). This dropout was caused by a substantial mismatch between our GRCh38-aligned scRNA-seq dataset and the pyscenic sourced feather file (motif-to-gene linking data) available online (https://resources.aertslab.org/cistarget/databases/; as of early 2026), despite the fact that the latter is also GRCh38-derived. This mismatch was caused by a combination of mismatching gene SYMBOLs in the feather file, and genes annotated in the scRNA-seq dataset that were absent in the feather file. Step 1 of pyscenic constructs Regulons based on transcription factor-(TF)-to-target gene expression correlation in the scRNA-seq data. Step 2 (Rcistarget) filters regulons on the basis of TF motif enrichment within 10kb of the putative target gene. The mismatch created two problems: (1) SYMBOL mismatch in TFs directly resulted in loss of regulons headed by those TFs. For example, the gene for the transcription factor TBXT/T/Brachyury was labelled as “TBXT” in our scRNA-seq dataset, but “T” in the feather file. For this reason, predicted Regulons for TBXT could not be matched to the correct motif information and so these regulons were dropped. (2) The Rcistarget pipeline drops any regulons for which fewer than 80% of targets are found in the feather file, with a warning message from pyscenic.transform saying the module will be skipped. Any targets within regulons that are present in the scRNA-seq dataset and absent in the feather file, or with mis-matched SYMBOLs, contribute to meeting this threshold. To fix this we generated a new feather file, using the R genome annotation package EnsDb.Hsapiens.v86 to fix mismatched SYMBOLs (∼600). We then created an omission list that we used to omit from the analysis any genes that were not present in the feather file before the Rcistarget step. Many of these remaining were adapter proteins. This resulted in the recovery of a large number of regulons. The updated feather file and the gene omission list are available upon request.

We also adapted the SCENIC pipeline to output negative regulons as well as positive. Negative (repressing) regulons can already be generated as part of steps1-2 of pyscenic (by using “all_modules = TRUE”) but step 3 (AUC regulon activity) is not compatible with repressing Regulons, as it ranks Regulons according to the cumulative expression of their targets. To address this, we used a custom python script (process_pyscenic_step2.py) to process the output of step 2, removing any rows for which the TargetGenes column was empty and then separating the positive and negative regulons into separate csv files. For the negative Regulons we then ran step 3 using a loom file with an inverted gene expression matrix (calculated per gene as maxCount-count), to allow ranking of repressing regulons.

SCENIC regulon-target networks were visualized in R using the tidygraph and ggraph packages, with edges weighted by SCENIC-derived target gene scores and styled to distinguish activating from repressing interactions. Where CUT&RUN data were incorporated, targets absent from the SCENIC output were overlaid as additional nodes connected by unweighted edges, with targets shared between both datasets visually distinguished from those exclusive to either source.

### ATAC-seq sample processing and analysis

#### Sample prep

ATAC-seq sample processing and sequencing was carried out by the IGO core at MSKCC. Profiling of chromatin was performed by ATAC-Seq as described previously ^188^. Briefly, 50,000 cells were washed in cold PBS and lysed. The transposition reaction was carried out using TDE1 Tagment DNA Enzyme (Illumina catalog # 20034198) incubated at 37°C for 30 minutes. The DNA was cleaned with the MinElute PCR Purification Kit (QIAGEN catalog# 28004) and material was amplified for 5 cycles using NEB Next High-Fidelity 2X PCR Master Mix (New England Biolabs catalog #M0541L). After evaluation by real-time PCR, 7-10 additional PCR cycles were done. The final product was cleaned by aMPure XPbeads (Beckman Coulter catalog # A63882) at a 1X ratio, and size selection was performed at a 0.5X ratio. Libraries were sequenced on a HiSeq 4000 in a PE100 run, using the HiSeq 3000/4000 SBS Kit (Illumina) as paired-end reads to a depth of 20–30 million reads per library.

#### Data analysis

ATAC-seq data was processed following the recommendations of the ENCODE consortium (The ENCODE Consortium ATAC-seqData Standards and Prototype Processing Pipeline https://www.encodeproject.org/atac-seq/). Raw sequencing reads were 3’ trimmed for both quality and adapter content using TrimGalore v0.4.5 and quality was evaluated with FastQC v0.11.5. Reads were aligned to human assembly hg38 with version 2.3.4.1 of bowtie2 using the default parameters and MarkDuplicates of Picard Tools version 2.18.6 was used for deduplication. samtools (v.1.9) was used to generate bam files and filter out low quality (Phred quality score <30) and mitochondrial reads. Enriched regions were determined using MACS2 v2.1.0 and a peak atlas was created from a superset of +/- 250 bp around peak summits from all samples. Differentially accessible regions (DARs) were called using DESeq2 (v.1.16)^189^ with pairwise comparisons (fold change ≥ 1.5, FDR ≤ 0.1). Normalized bigwigs files were generated from bams using deepTools (v.3.5.1) bamCoverage (--scaleFactor 10 --normalizeUsing CPM --extendReads, -- ignoreDuplicates) and averaged across replicates per condition using bigwigAverage. Tracks visualized in IGV and heatmaps were generated from averaged bigwigs. For heatmaps “bump” vs. “no bump” enriched DARs are presented for all samples and were generated using deepTools computeMatrix (reference-point, centered on the peak midpoint, ±3,000 bp) and plotHeatmap. Pathway enrichment was performed on genes associated with differential peaks using the enrichGO function in the Bioconductor package clusterProfiler^189^. Transcription factor footprinting was performed using TOBIAS (v.0.13.3). ATACorrect and ScoreBigwig were run on each replicate BAM/peak file, and BINDetect was used to identify differentially bound motifs between conditions from averaged footprint-score bigwigs (replicate 2 was excluded due to poor signal-to-noise) against the merged peak atlas, using the HOCOMOCO (H14 CORE) motif database (motif p-value 1×10⁻³, bound probability threshold 0.0001).

### CUT&RUN sample processing and analysis

#### Sample prep

Cleavage Under Targets & Release Using Nuclease (CUT&RUN) was performed using CUTANA ChIC/CUT&RUN kit (Epicypher #14-1048) following an optimized version of manufacturer’s protocol. Briefly, 1 × 10^6^cells were coupled with Concanavalin A beads, permeabilized with Digitonin at a final concentration of 0.05% and mixed with 0.5 μg target antibody in antibody buffer (antibodies are listed in Table 1.). 2μL of SNAP-CUTANA™ K-MetStat Panel were added to reactions designated for histone modification antibodies. All samples were incubated overnight on a nutator at 4C. The following day, cells were first incubated for 10 minutes with micrococcal nuclease fused to proteins A and G (pAG-MNase), which was then activated by CaCl_2_ addition. After 2h incubation at 4C, the reaction was stopped with stop buffer. DNA was then isolated using SPRI beads and quantified with Qubit. Libraries were prepared using NEBNext Ultra II DNA library kit, following manufacturer’s recommendations. Fragment size was detected using a Tape Station system and multiplexed libraries were sequenced on a NExtSeq2000 instrument (Illumina) at 50bp paired end reads.

#### Data analysis

Adapters were trimmed using NGmerge (v.0.3) for paired-end reads. Reads were aligned to the human reference genome (GRCh38/hg38) using Bowtie2 (v.2.4.1) with parameters --very-sensitive-local, --no-mixed, --no-unal, --no-discordant, and read quality was assessed using fastQC. samtools (v.1.9) was used to generate bam files and filter out low quality (Phred quality score <30), mitochondrial and duplicated reads. Significant binding peaks were determined using macs2 v2.1.0 for narrow peaks, while –broad parameter was added for broad peak analysis. Negative IgG files were used as control in peak calling with two biological replicates per condition. Cut-off values (q-values) were determined post-hoc, testing several q-values based on signal-to-background ratio and using macs2 -- cutoff-analysis parameter for additional guidance. Peaks located in ENCODE blacklisted regions were excluded. Fraction of Reads in Peak (FRiP) scores were calculated as additional quality control. Coverage tracks (bigwig files) were generated from filtered bam files using deepTools (v. 3.5.1) bamCoverage with parameters --normalizeUsing RPGC --binsize 1 and visualized in IGV alongside ATAC-seq accessibility tracks. Metagene plots and heatmaps showing signal enrichment over genomic regions were generated by first calculating intensity scores for each genomic site using deepTools computeMatrix and then plotting heatmaps and/or metagene plots with plotHeatmap, selecting the center of the peaks as reference point. For final peak representation, bam files from different replicates were merged into a master file using samtools merge before generating coverage tracks and heatmaps. For differential binding analysis, CUT&RUN data from two different biological replicates were analyzed using DiffBind (v. 3.4.11), along with matching IgG as control, following the package documentation (http://bioconductor.org/packages/release/bioc/vignettes/DiffBind/inst/doc/DiffBind.pdf). Briefly, a binding matrix was generated with dba.counts (summits=250). Count normalization was performed using the DBA_NORM_LIB parameter. DESeq2 was used to perform differential peak analysis.

#### Histone-state analysis

For DARs identified in the “bump” vs. “no bump” ATAC-seq analysis, mean CUT&RUN signal for H3K4me3 and H3K27me3 was extracted over a 500 bp window centered on the nearest TSS using deepTools multiBigwigSummary. Signal was quantified from RPGC-normalised (1× genome coverage) tracks, so that values are expressed in units of genome-average coverage, and log2-transformed following addition of a pseudocount of 1 — equivalent to one genome-average of background, and matched to the scale of the data so that regions lacking a mark are not displaced from the measured distribution. A contrast score, log2((H3K4me3 + 1) / (H3K27me3 + 1)), and a magnitude score, log2(H3K4me3 + 1) + log2(H3K27me3 + 1), were calculated per region. Because RPGC normalisation places both marks on a common coverage scale, no further per-mark scaling was applied: the contrast score is therefore a direct log2 fold-difference between the two marks, and the magnitude score a measure of total histone signal, with the origin corresponding to absence of both marks. Plotting magnitude against contrast is a 45° rotation of the log–log scatter of the two marks, analogous to an MA (Bland–Altman) plot, separating the overall level of histone marking from the direction in which it is skewed.

Regions were classified as Putative Active (contrast > 1, i.e. ≥ 2-fold H3K4me3 dominance over H3K27me3), Putative Inactive (contrast < −1) or Putative Poised (|contrast| ≤ 1) chromatin signatures, with regions of magnitude < 1 classified as Ambiguous. The magnitude threshold was set equal to the contrast threshold since a region carrying only one mark has magnitude equal to |contrast|, this is the lowest floor that excludes single-marked regions from the Putative Poised class, while imposing no additional requirement on the signal a lone mark must reach to be classified.

### Calcium imaging

Changes in intracellular calcium concentration ([Ca²⁺]ᵢ) were measured by Fluo-4 AM fluorescence imaging, as described previously ^56,85^. Cells were loaded with 5µM Fluo-4 AM (Invitrogen) for 20 min at room temperature in HEPES-buffered physiological saline (149mM NaCl, 5mM KCl, 1mM MgCl₂, 2mM CaCl₂, 10mM glucose and 10mM HEPES) and subsequently washed for 20 min in dye-free HEPES solution. Recordings were acquired for 2 min, including a 10s baseline period followed by local application of agonists diluted in HEPES solution (high-potassium solution for 5s [78mM NaCl, 75mM KCl, 10mM HEPES, 2mM CaCl₂, 1mM MgCl₂ and 10mM D-glucose], 20µM 1,1-dimethyl-4-phenylpiperazinium [DMPP; Fluka] for 10 s, 1µM Substance P [Sigma] for 10s, or 10µM 5-HT [Sigma] for 10 s). Each stimulus recording was followed by a 3-min washout with HEPES solution. Image sequences were analysed in Igor Pro (WaveMetrics) using custom-written macros developed by Pieter Vanden Berghe (KU Leuven)^190^. Regions of interest were manually drawn around the cell body of each responding cell, and fluorescence intensity (F) was extracted for each region. Fluorescence values were normalised to baseline fluorescence (F₀), calculated as the mean fluorescence intensity of the first 10 frames, and responses were expressed as F/F₀.

Calcium responses were included only when the increase in fluorescence exceeded intrinsic signal noise by at least tenfold. Response amplitude was defined as the maximum F/F₀ value above baseline. Response duration was quantified as the time required for the calcium signal to decay from its peak to 50% of the peak amplitude.

### RNA isolation and qPCR

Cells were collected in Trizol and processed using Direct-zol RNA Mini-prep kit (Zymo Research, R2050). A total of 0.5μg of RNA was used to generate cDNA using iScript (Bio-Rad). qPCR was performed using SsoFAST EvaGreen Mix (Bio-Rad) in a Bio-Rad CFX96 Thermal Cycler. The manufacturers protocol was used for all steps.

### Statistics and reproducibility

Sample sizes were estimated based on previous publications in the field. Investigators were not blinded to experimental conditions. Transcriptional and genomic studies were performed with the same bioinformatic pipeline between conditions. Statistics and data representation were performed using either R or PRISM (GraphPad) unless otherwise specified in the methods. Statistical tests used for each analysis are indicated in the figure legend. Data are represented as mean ± s.e.m. unless otherwise indicated.

## Acknowledgements

The authors thank members of the Studer laboratory for discussion and insights into the project; M. Lisi and A. Pepe-Caprio for assistance and laboratory management; Samantha Morris and Wenjun Kong for CellTag support; Ronan Chaligné for CellTag experimental design input; the Stem Cell Core at MSKCC (RRID: SCR_027808) for their support with CRISPRa and generation of reporter cell lines; the Gene Editing & Screening Core at MSKCC (RRID:SCR_022606) for their support with CellTag libraries; the Center for Epigenetics Research at MSKCC (RRID:SCR_022519) for their support with ATAC-seq anaylsis and epigenomics advice; the Integrated Genomics Operation at MSKCC (RRID:SCR_022523) for their support with sequencing; the Single Cell Analytics Innovation Lab at MSKCC (RRID:SCR_022527) for their support with sequencing and genomics experimental design; the Molecular Cytology core at MSKCC (RRID:SCR_022525) for their support with imaging. This work was supported by NINDS/NIH (R01NS099270; R01NS118067), NICHD/NIH (R21HD112917), NIH/NCI Cancer Center Support Grant (P30 CA0088748; GES core), the MSKCC-Therapeutic Development Fund, The Michael J Fox Foundation (MJFF-026800), the Freedom Together Foundation (FTF), a Parkinson’s Foundation Launch Award (PF-LAUNCH-1584415), a Horizon MSCA fellowship (HORIZON-MSCA-2024-PF-01: 101203036), a sponsored research agreement from BlueRock Therapeutics (SK2018-1304) and a grant from the Australian NHMRC MRFF Stem Cell Therapies Mission (2032801).

## Author contributions

J.O.S.H. and L.S. conceived of the project, interpreted the results and wrote the manuscript. J.O.S.H. developed, performed and interpreted all wet lab experiments and data analysis unless otherwise stated. G.A. performed and co-developed the 16-condition “binary approach” experiment. R.V. performed and co-developed the mDA neuron experiments with the 6-cell line pool and H9-GBX2::tdTOM. X.W. performed the validation of the bump protocol in H1 and KOLF. V.D.B. performed CellTypist analysis as well as performed and co-developed the SCENIC analysis approach and fix. E.S.R. performed the calcium imaging and analysis. D.Y. performed mDA neuron experiments in KOLF and H9-PITX3::GFP lines. Y.W. designed CRISPRa guides, advised on this experiment and performed the electroporation. S.M. performed the CellTag promoter swap and made the libraries. A.Z. constructed the H9-GBX2::tdTOM reporter line. E.G. and V.CK. performed and advised on the CUT&RUN. R.P.K. performed ATAC-seq analysis up to the differential enrichment analysis and advised on epigenetic approach. R.M.W. made the GFP-H2B, tdTOM-H2B and 3xFLAG-H2B cell lines used in the assembloid experiment. A.E. aided in the mDA neuron experiments. Y.F. contributed to early conceptual framework. E.B. supervised CUT&RUN. R.J.G. supervised CellTag library construction. L.A.S. and M.M.H. supervised the calcium imaging. T.Z. supervised the CRISPRa approach and H9-GBX2::tdTOM reporter cell line construction.

