## Supplementary Figures for "Timed binary WNT activity is sufficient to model the early human neural axis"

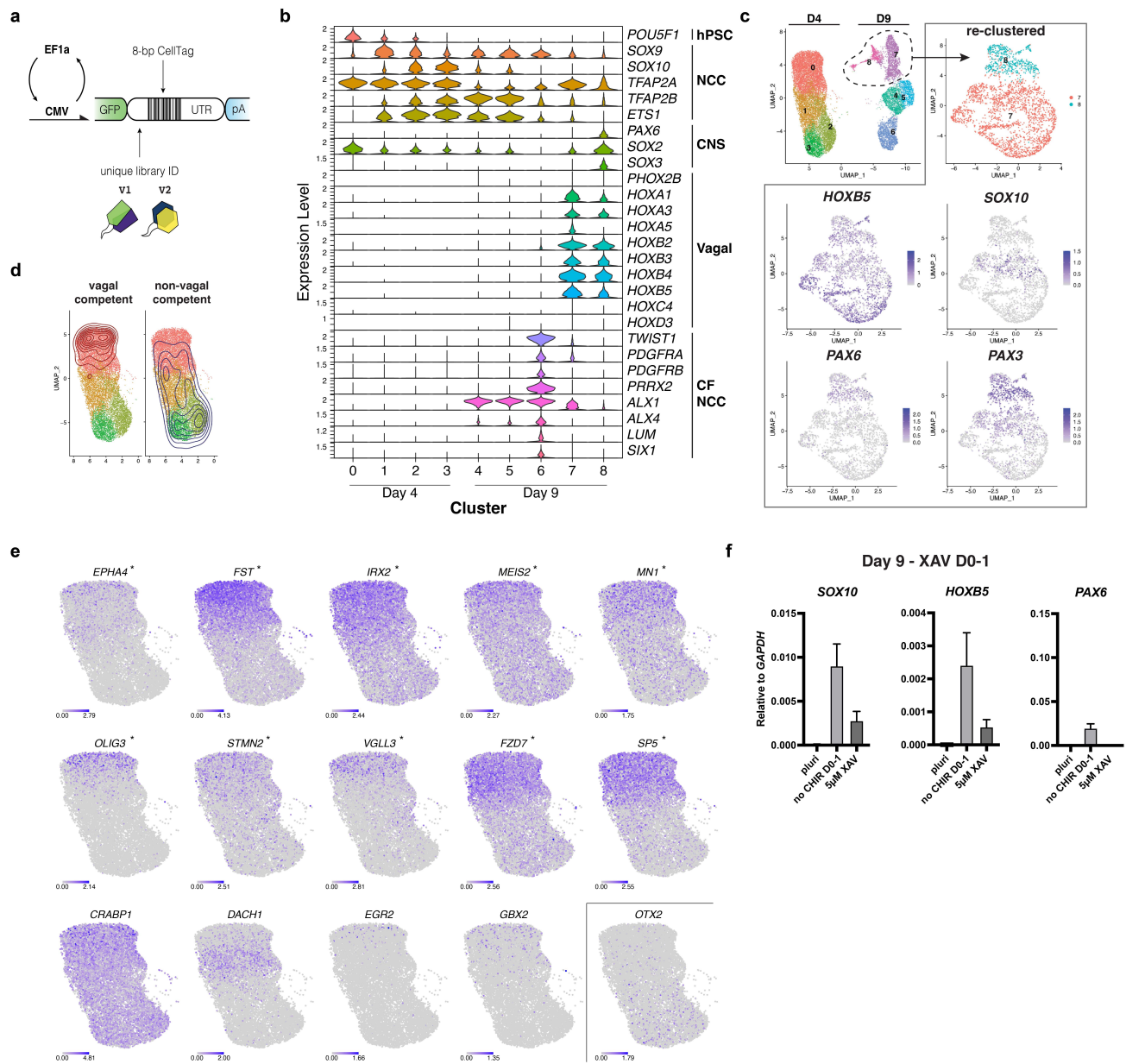

**Supplementary Figure 1 - Vagal competent clones exhibit hindbrain-level transcriptional features and generate both CNS and NC derivatives. (a)** Schematic of CellTag transgene showing promoter exchange. **(b)** Violin plots showing expression of genes of interest from CellTag scRNA-seq data. CF = cranio-facial **(c)** Sub-clustering of vagal clusters shows both neural crest and CNS lineages. **(d)** Location of cells belonging to vagal and non-vagal competent clones at Day 4 only. This gives a better representation of distribution of cells than when Day 9 is also present. **(e)** Gene overlays for day 4 showing genes of interest enriched in vagal competent clones (\*) and enriched in cluster "0" (bottom row except *OTX2*). *OTX2* is shown as an anterior marker for contrast **(f)** Day 9 - qPCR after inhibition of WNT signaling D0-1 using XAV939. XAV treatment reduces the proportion of spontaneous vagal competent cells but also reduces neural crest differentiation efficiency. Interpreted as anteriorization to forebrain level where NCC do not form.

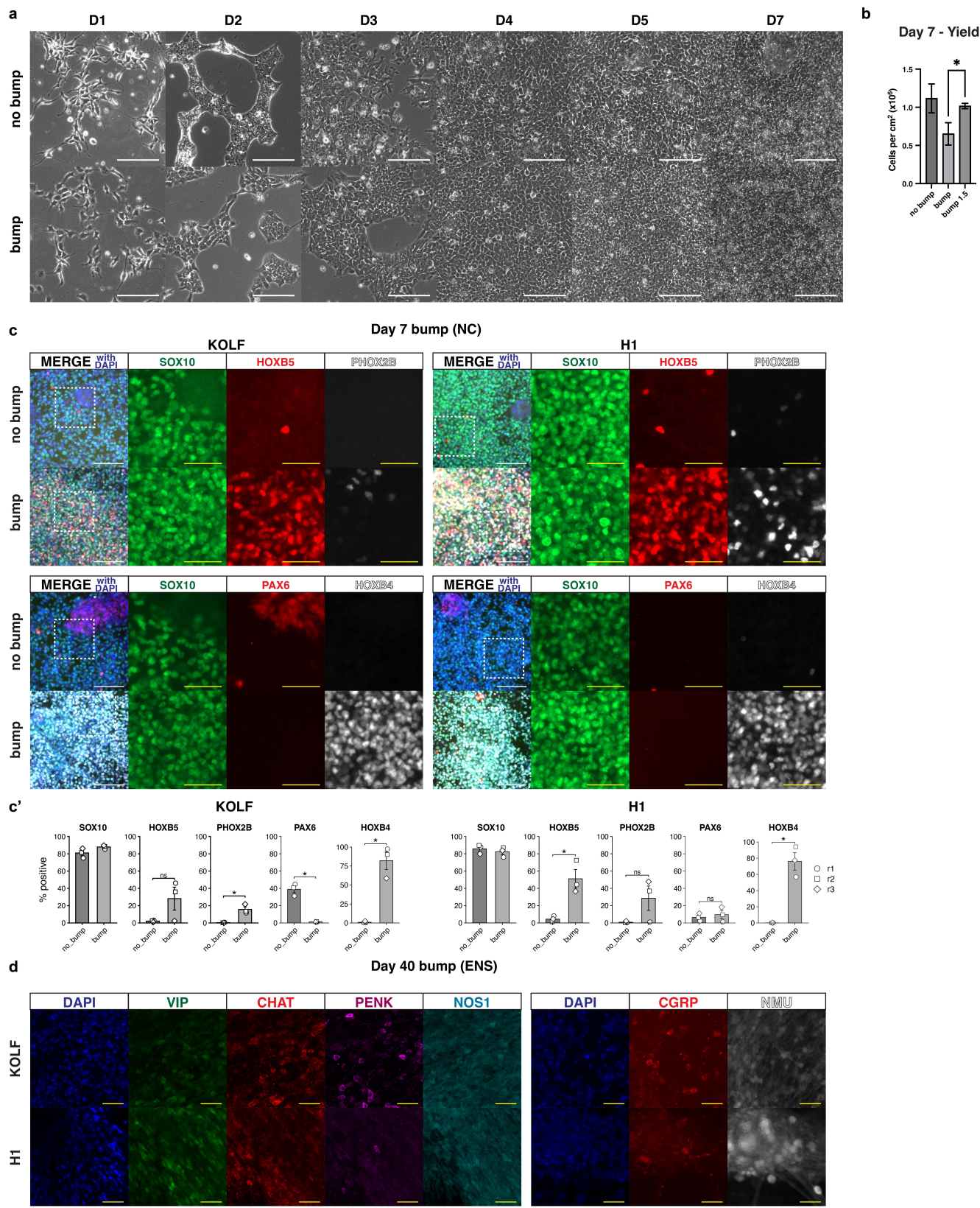

**Supplementary Figure 2 - Early, strong, transient WNT activation induces vagal competence in multiple cell lines. (a)** Phase-contrast images showing culture morphology during neural crest differentiation with “no bump” and “bump” and “bump 1.5” D4-7 cultures using H9-*SOX10::GFP*. **(b)** Day 7 cell yield per cm<sup>2</sup>. “Bump 1.5” represents condition where CHIR concentration is increased to 1.5μM from D4-7 with improved yield relative to “bump”. **(c)** ICC demonstrating upregulation of vagal markers HOXB5, PHOX2B and HOXB4 at day 7 of NC induction using the WNT bump protocol in hiPSC line KOLF and hESC line H1. **(c')** Quantification of (c). **(d)** ENS D40 ICC showing VIP, CHAT, NOS, PENK, CGRP and NMU in KOLF and H1. ICC quantification data in (c') are presented as mean ± s.e.m. Each point represents a biological replicate. Two-tailed unpaired t-tests were used with Welch's correction for unequal variances. \*P < 0.05, \*\*P < 0.01, \*\*\*P < 0.001; ns, not significant (P ≥ 0.05). Scale bar: White - 100μm; yellow - 50μm.

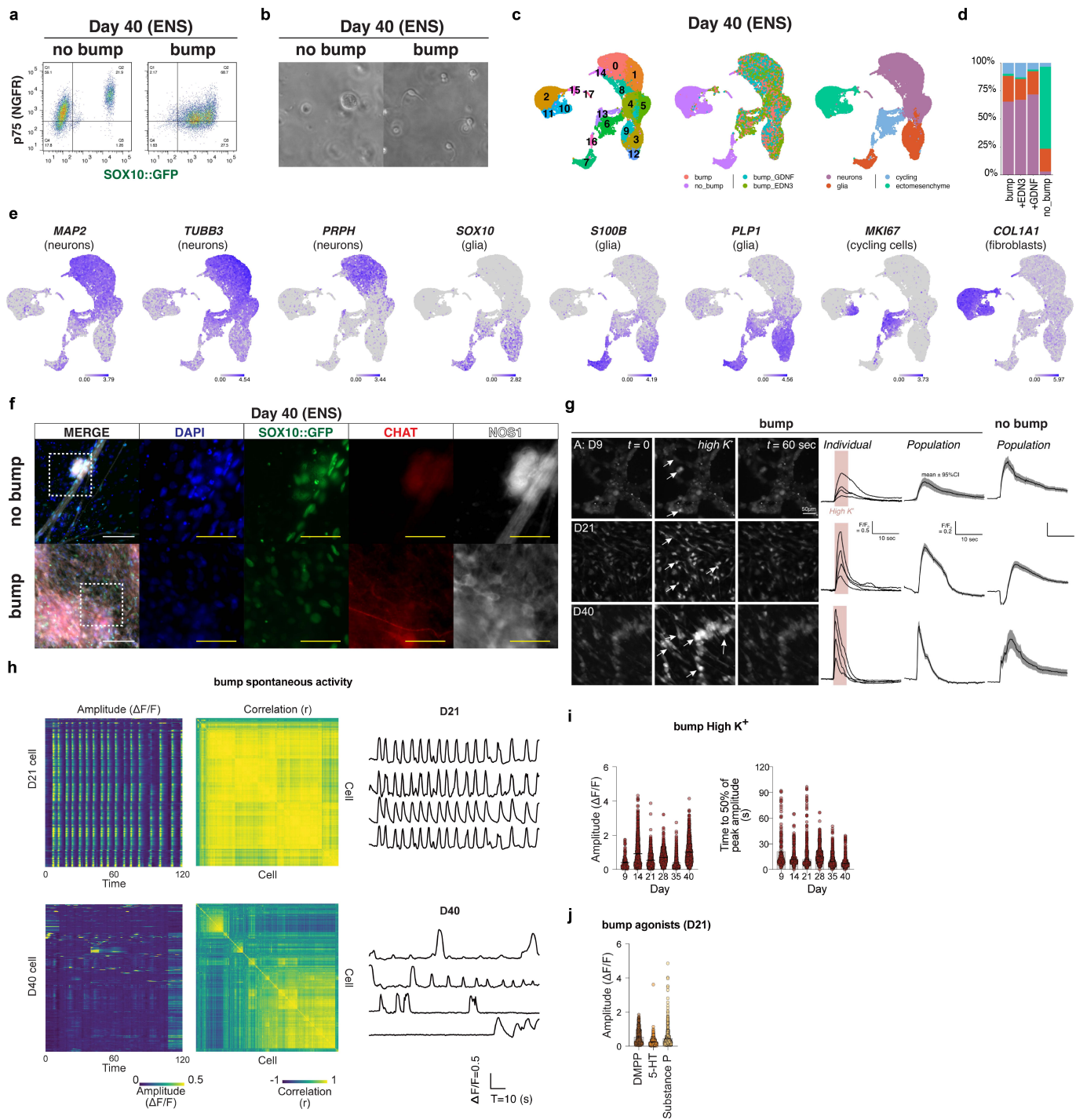

**Supplementary Figure 3 - Improved vagal patterning eliminates off-target ectomesenchymal lineages in VNC-derived ENS cultures and results in faster maturation of neurons.** (a) ENS day 40 flow cytometry showing p75 and SOX10::GFP detection post-dissociation. (b) ENS day 40 phase-contrast imaging of single cell suspension of dissociated ENS cultures prior to flow cytometry and sequencing. (c) UMAPs representing scRNA-seq analysis of day 40 ENS cultures derived from WNT “bump” or “no bump” cultures. “bump\_EDN3” and “bump\_GDNF” conditions represent variants of the “bump” protocol where EDN3 (100nM) or GDNF (25ng/mL) were added to the media between D4-7. (d) Stacked bar chart representing contribution to distinct lineage types across the different samples. Ectomesenchymal cell types make up 75% of “no bump” cultures but ~2% of “bump” cultures. (e) Gene expression overlays on UMAP in (c) for major lineage markers. (f) Day 40 ENS ICC using H9-SOX10::GFP and staining for NOS1 and CHAT. Neurons still exhibit SOX10::GFP signal, albeit lower than in glia, presumably because turnover of GFP-H2B in post-mitotic cells is slow. Transcripts for SOX10 are not present in neurons (see e). (g) Fluorescence images of Fluo-4 AM loaded cells tracing Ca<sup>2+</sup> response at day 9 (1st row), day 21 (2nd row) and day 40 (3rd row) at baseline (T=0), following high K<sup>+</sup> stimulation (high K<sup>+</sup>), and post-transient (T=60) in the WNT “bump” condition. Active cells are marked with arrows. Representative individual and population-level traces are shown to the right of each panel alongside a population trace for the WNT “no bump” condition at the same time point. (h) Heatmap and representative traces of spontaneous activity in WNT “bump” ENS cultures at day 21 and day 40. Early synchronous spontaneous activity at day 21, a hallmark of early neural development, gives way to activity in discrete networks by day 40, a hallmark of mature neurons. (i) Amplitude and duration of [Ca<sup>2+</sup>]<sub>i</sub> transients in “bump”-derived ENS cultures over time showing all time points tested. (j) Amplitude of [Ca<sup>2+</sup>]<sub>i</sub> response to the nicotinic acetylcholine receptor (nAChR) agonist DMPP, the NK1 receptor agonist Substance P, and serotonin (5-HT) at day 21 in the WNT “bump” condition. Scale bar: White = 100µm; yellow = 50µm.

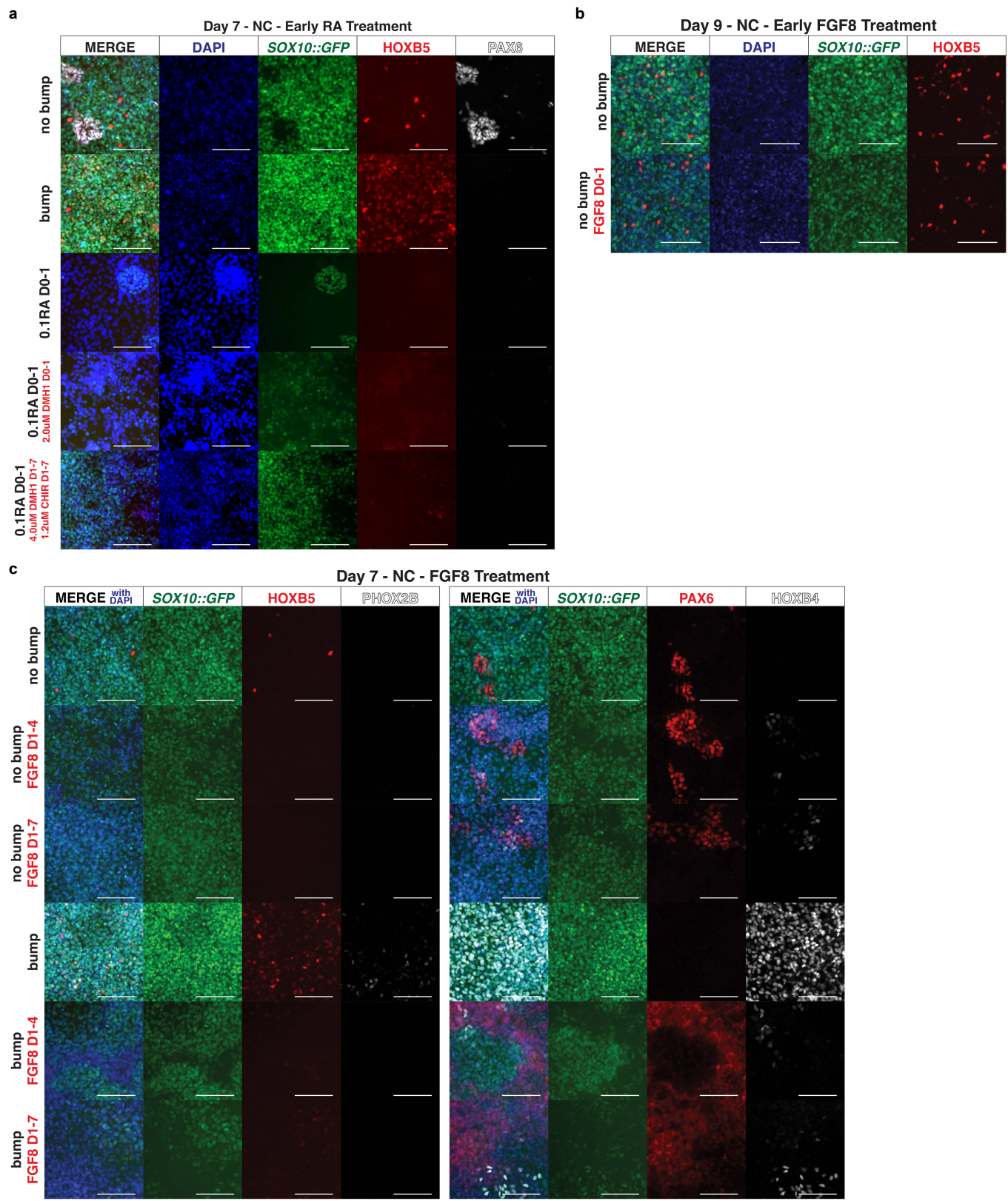

**Supplementary Figure 4 - Early treatment with retinoic acid or FGF8 does not induce vagal competence.** (a) Day 7 NC ICC after activation of RA signaling between days 0-1 instead of CHIR. Early RA treatment alone induces a non-neural ectoderm-like phenotype. This can be accounted for by treatment with 2.0μM DMH1 from days 0-1 (row 4) or by treatment with 4.0μM DMH1 and 1.2μM CHIR from days 1-7 (row 5), but the resulting neural crest do not show expression of vagal markers HOXB5 and PHOX2B. (b) Day 9 NC (using protocol described in Figure 1d) ICC after treatment with FGF8 from days 0-1. No increase in the proportion of vagal NCC is observed. (c) Day 7 NC ICC after FGF8 treatment from days 1-4 or days 1-7. No increase in the proportion of vagal NCC is observed in the no bump condition and expression of vagal markers is reduced in the bump condition alongside a reduction in neural crest induction efficiency. Scale bar - 100μm.

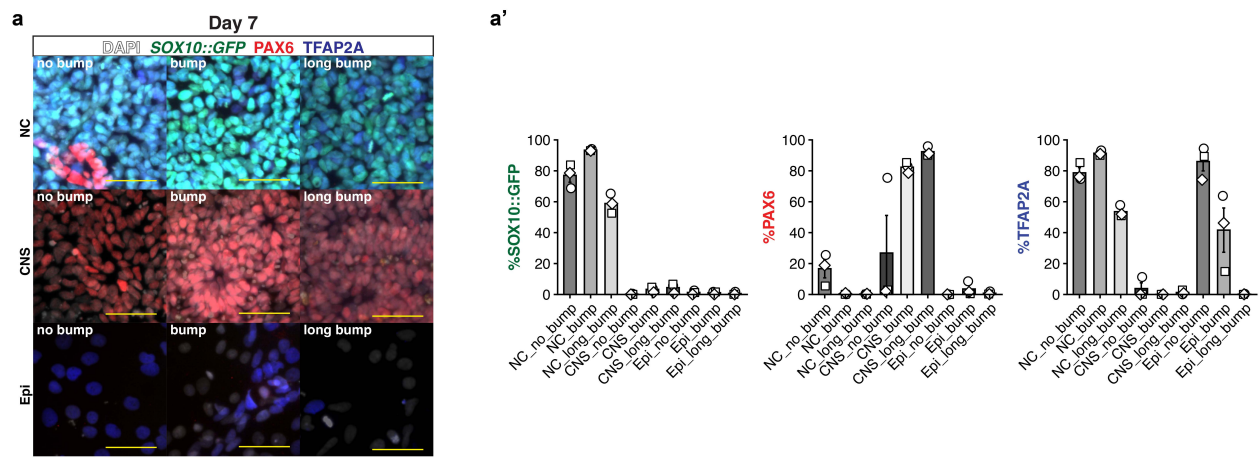

**Supplementary Figure 5 - Axially committed progenitor populations exhibit pan-ectodermal potential. (a)** ICC showing behaviour of “no bump”, “bump”, and “long bump” progenitors in neural crest (NC; intermediate BMP activation), central nervous system (CNS; BMP inhibition) and Epidermal (Epi; strong BMP activation) induction conditions with quantification in (a'). Scale bar - 50µm.

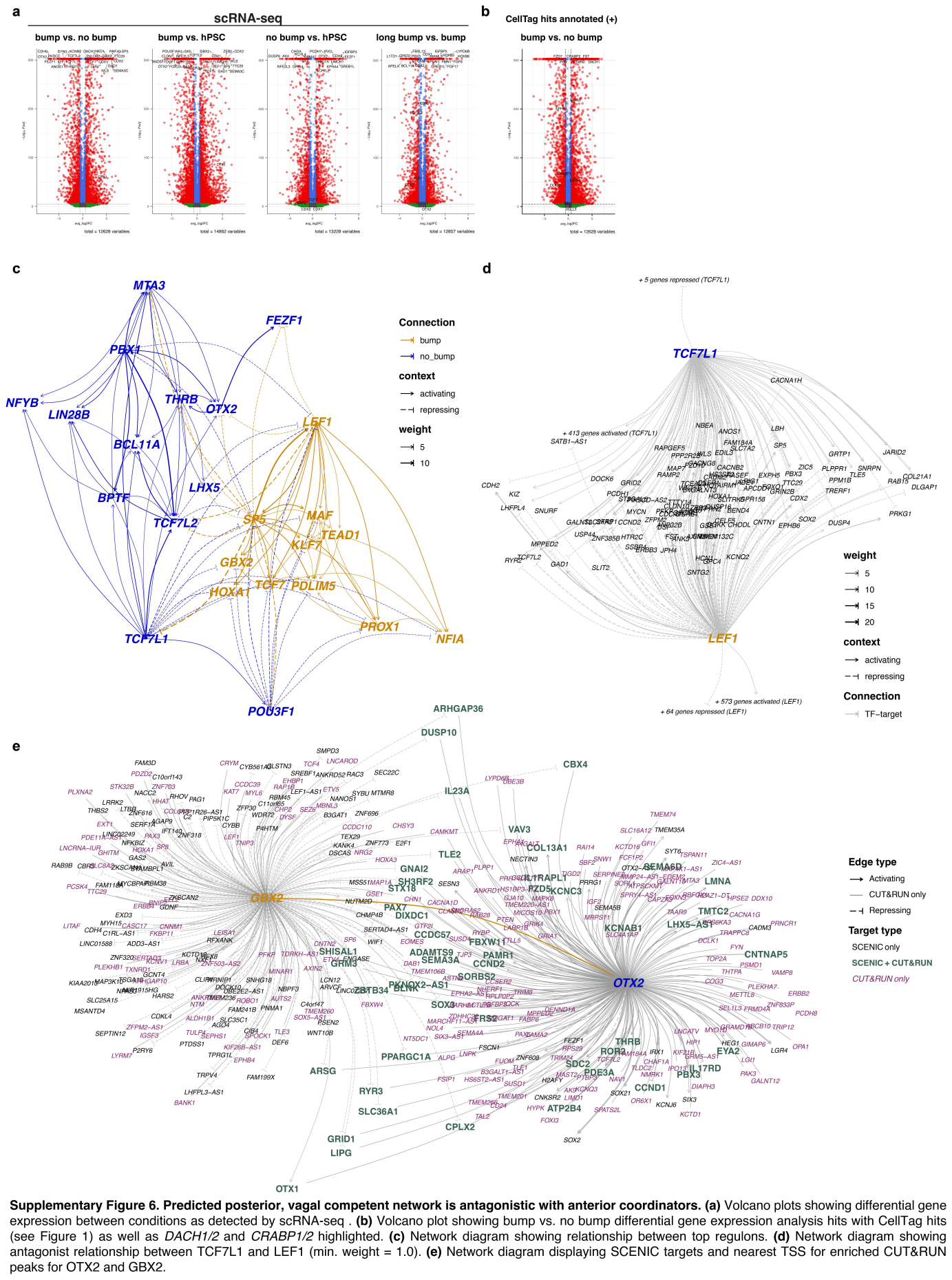

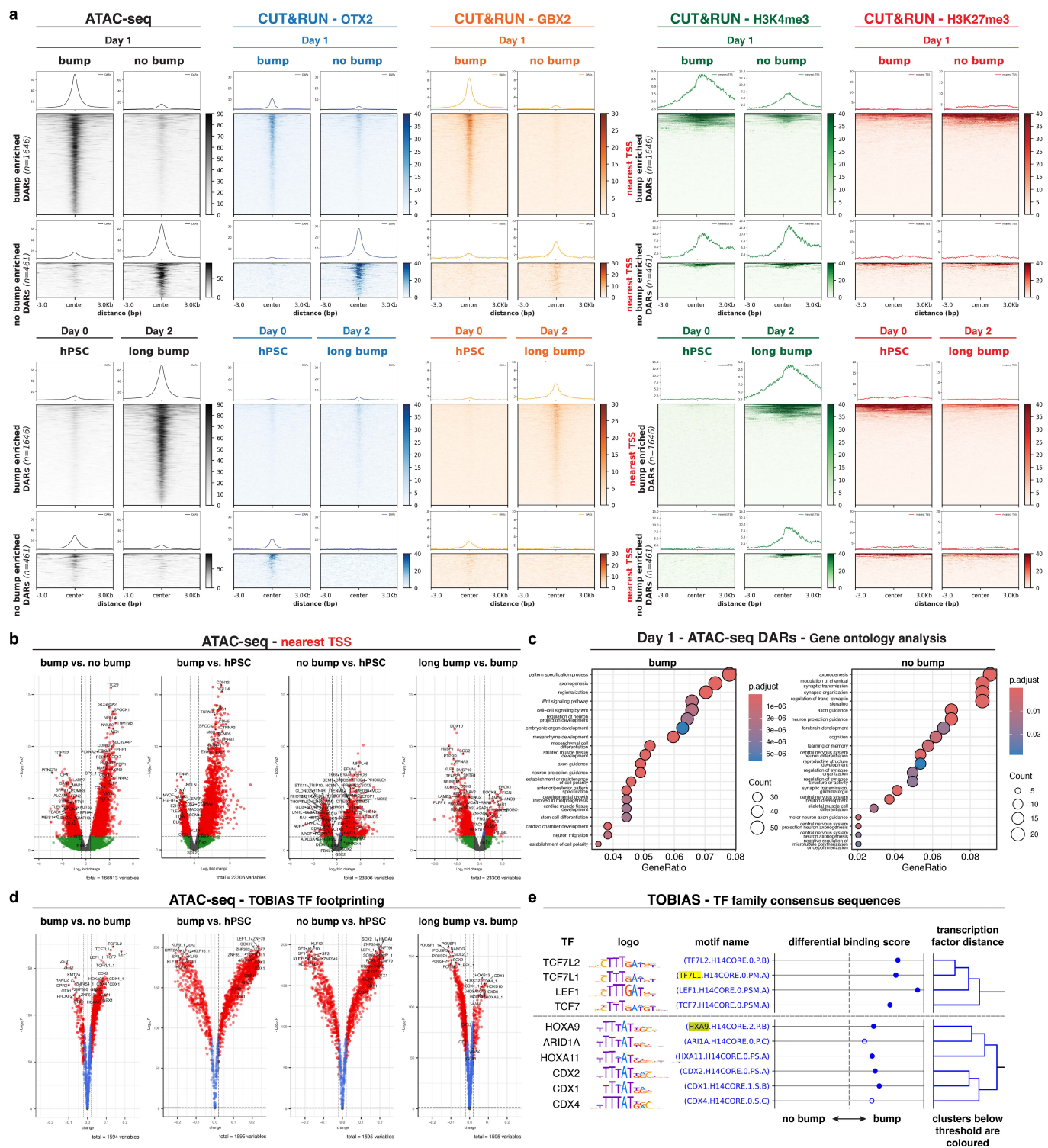

**Supplementary Figure 7. Evidence of axial fate coordinator binding in differentially accessible regions at day 1. (a)** Heatmaps showing ATAC-seq and CUT&RUN signal across differentially accessible regions (DARs; accessibility, OTX2, GBX2) or nearest transcriptional start site to DARs (TSS; H3K4me3, H3K27me3). Data are shown for “bump” & “no bump” (top) and hPSC & “long bump” (bottom). **(b)** Volcano plots showing significantly enriched and depleted DARs across different pairwise comparisons with nearest TSS shown for top hits. **(c)** Gene ontology analysis for “bump” vs. “no bump” DARs shown in (b). **(d)** Volcano plots showing differential transcription factor (TF) footprinting analysis using TOBIAS. **(e)** TF family binding site similarity - output from TOBIAS. Motifs clustered by similarity. Coloured subclusters indicate motifs are too similar to distinguish with confidence. Filled circle for differential binding score indicates significance.

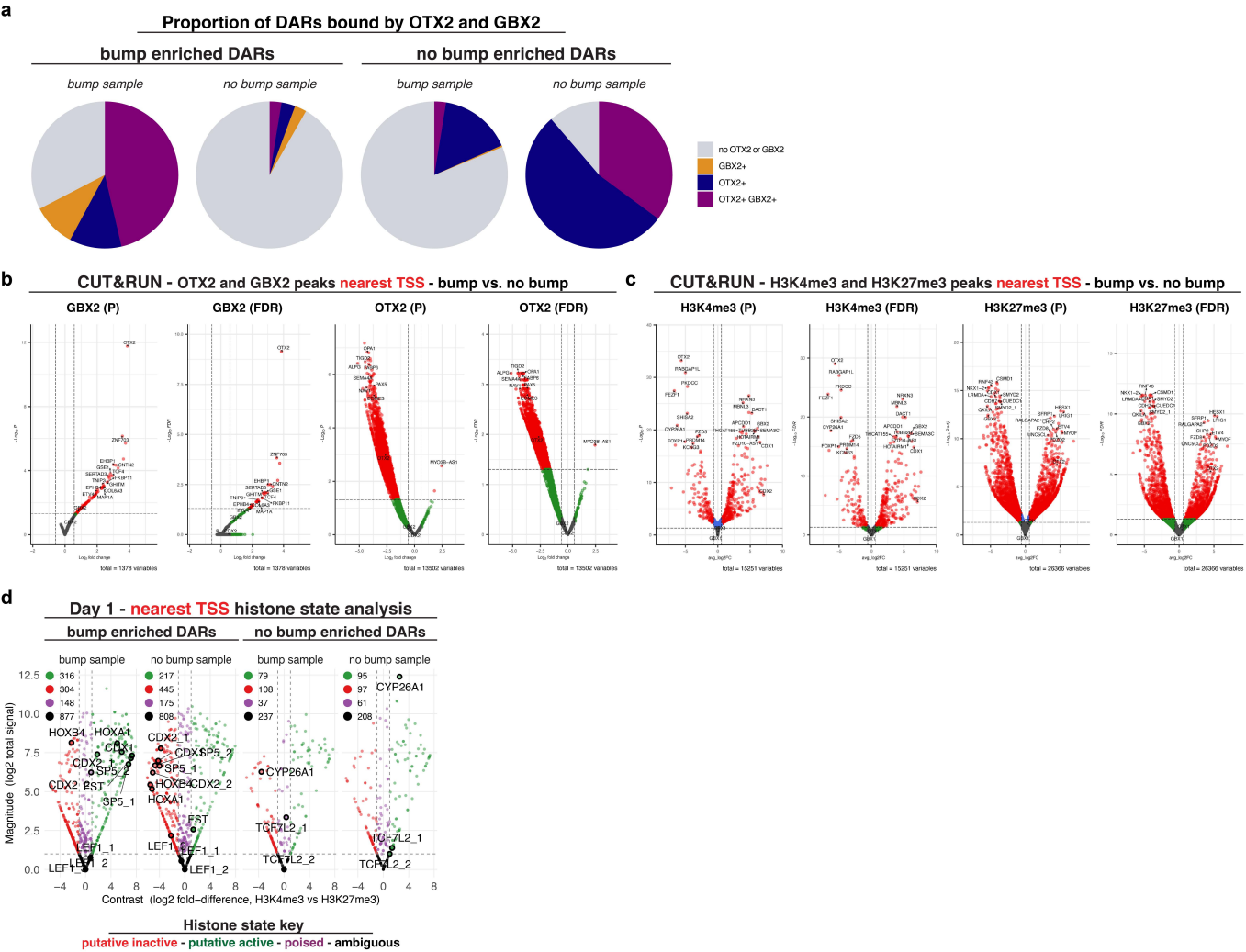

**Supplementary Figure 8. Axial fate coordinators are directly bound by OTX2 and GBX2 and exhibit distinct histone signature between “bump” and “no bump” conditions. (a)** Proportion of ATAC-seq-derived DARs bound by OTX2 or GBX2 (CUT&RUN analysis). **(b)** Volcano plots showing differential binding of GBX2 and OTX2 for bump vs no bump with nearest TSS shown for top hits. Plots are shown for both p value and FDR. **(c)** Volcano plots showing differential binding of the activation marker H3K4me3 and the repression marker H3K27me3 for “bump” vs “no bump” with nearest TSS shown for top hits. Plots are shown for both p value and FDR. **(d)** Histone state analysis summarizing H3K4me3 and H3K27me3 CUT&RUN data for the nearest TSS to enriched DARs from the ATAC-seq data. Analysis annotates as putative active (high H3K4me3; green; right of plot), inactive (high H3K27me3; red; left of plot) poised (high for both marks; purple; middle of plot) or ambiguous (low for both marks; black; bottom of plot). Number of TSS on each plot is recorded at the top left of the plot. TSSs for multiple genes of interest are highlighted.

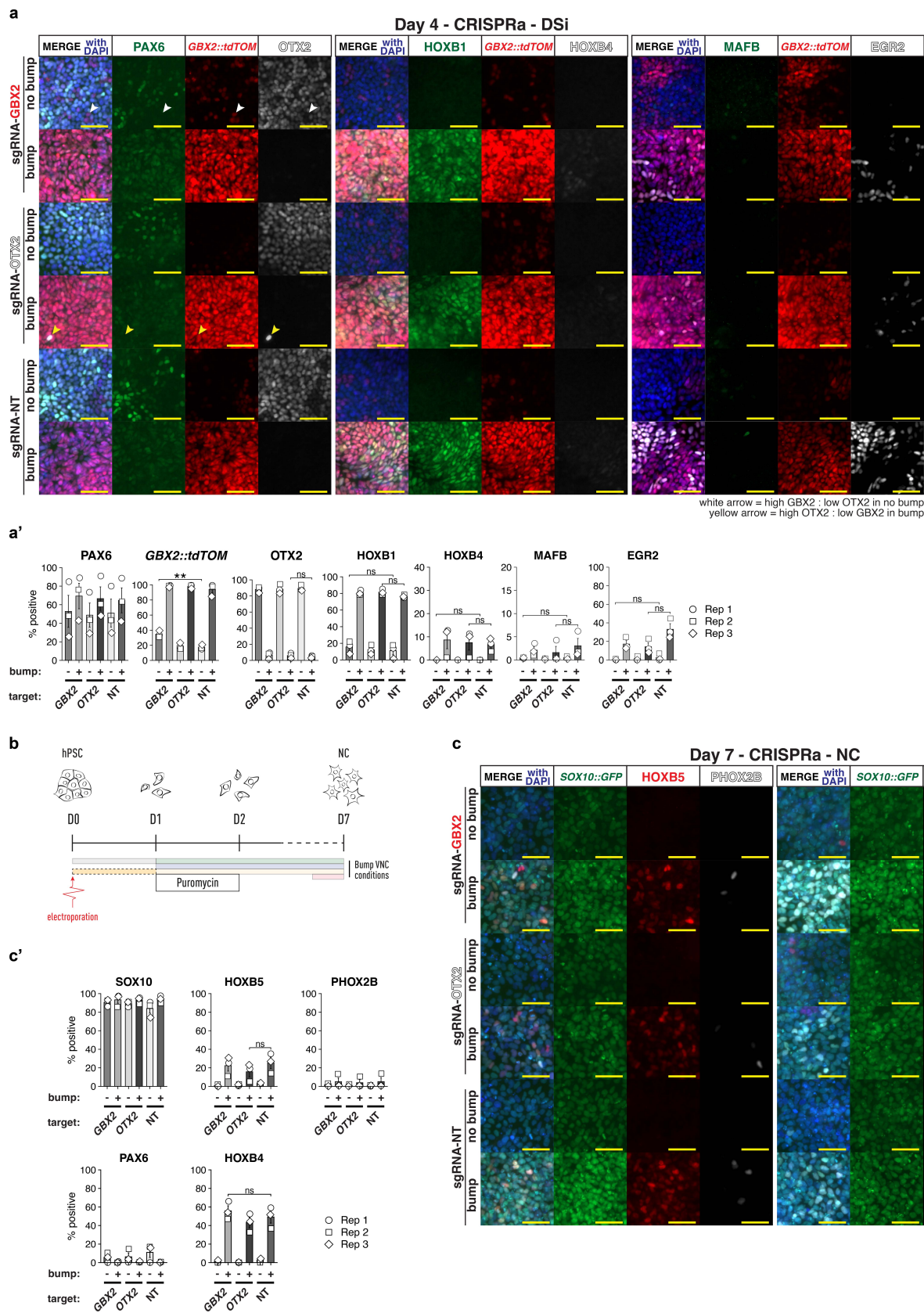

**Supplementary Figure 9 - CRISPRa-based activation of *OTX2/GBX2* alone does not affect long term axial patterning in CNS or neural crest cultures. (a)** ICC on day 4 of dual SMAD inhibition (DSi) after activation of *GBX2* or *OTX2* in “bump” and “no bump” conditions. Quantification of ICC in (a'). **(b)** Schematic showing experimental design of CRISPRa experiments in neural crest induction conditions. **(c)** ICC on day 7 of vagal neural crest induction after activation of *GBX2* or *OTX2* in “bump” and “no bump” conditions. Quantification of ICC in (c'). ICC quantification data in (a', c') are presented as mean ± s.e.m. Each point represents a biological replicate. Two-tailed unpaired t-tests were used with Welch's correction for unequal variances. \*$P < 0.05$, \*\*$P < 0.01$, \*\*\*$P < 0.001$; ns, not significant ($P \geq 0.05$). Scale bar - 50μm.

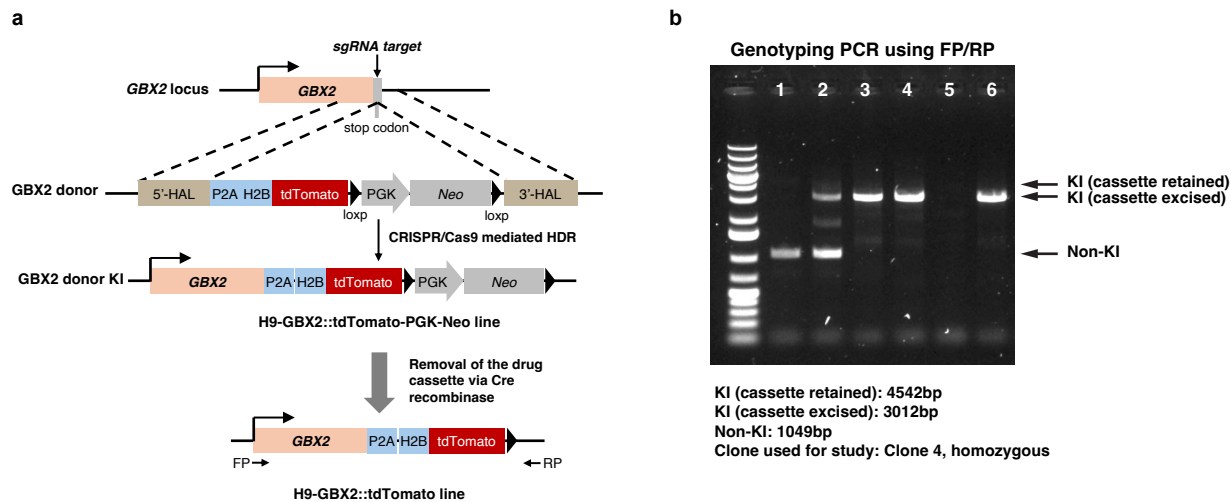

**Supplementary Figure 10 - Generation of the H9-*GBX2::tdTomato* reporter line.** (a) Schematic illustrating the generation of the H9-*GBX2::tdTomato* reporter line. (b) PCR verification of knock-in (KI) clones following Cre-mediated excision of the drug-selection cassette.
